# Interaction between a J-domain co-chaperone and a specific Argonaute protein contributes to microRNA function in animals

**DOI:** 10.1101/2024.02.27.581607

**Authors:** Pierre-Marc Frédérick, Guillaume Jannot, Isabelle Banville, Martin J. Simard

## Abstract

MicroRNAs (miRNAs) are essential regulators of several biological processes. They are loaded onto Argonaute (AGO) proteins to achieve their repressive function, forming the microRNA-Induced Silencing Complex (miRISC). While several AGO proteins are expressed in plants and animals, it is still unclear why specific AGOs are strictly binding miRNAs. Here, we identified the co-chaperone DNJ-12 as a new interactor of ALG-1, one of the two major miRNA-specific AGOs in *Caenorhabditis elegans*. DNJ-12 does not interact with ALG-2, the other major miRNA-specific AGO, and PRG-1 and RDE-1, two AGOs involved in other small RNA pathways, making it a specific actor in ALG-1-dependent miRNA-mediated gene silencing. The loss of DNJ-12 causes developmental defects associated with defective miRNA function. Using the Auxin Inducible Degron (AID) system, a powerful tool to acutely degrade proteins in specific tissues, we show that DNJ-12 depletion hampers ALG-1 interaction with HSP70, a chaperone required for miRISC loading *in vitro*. Moreover, DNJ-12 depletion leads to the decrease of several miRNAs and prevents their loading onto ALG-1. This study uncovers the importance of a co-chaperone for the miRNA function *in vivo* and provides insights to explain how different small RNAs associate with specific AGO in animals.

## INTRODUCTION

MicroRNAs (miRNAs) are endogenous small RNAs that regulate gene expression through base pairing with complementary mRNA targets (1, 2). These small noncoding RNAs possess a high regulatory potential since they can bind multiple mRNA targets, and one single mRNA can be bound by several different miRNAs. miRNAs are initially transcribed in the nucleus by the RNA polymerase II as primary miRNAs (pri-miRNAs). Pri-miRNAs are processed by the microprocessor formed by DGCR8 and Drosha to form precursor miRNAs (pre-miRNAs), which are exported to the cytoplasm. The pre-miRNAs are then processed by the RNAse III-type enzyme Dicer to form mature miRNA duplexes. After, the miRNA duplex is loaded onto Argonaute (AGO), and only one strand (called the guide strand) is selected to form the miRISC, the active silencing complex (3).

AGOs are bi-lobed proteins required for small RNA-mediated post-transcriptional gene silencing (4). They are composed of six domains. The two domains of the first lobe, the N-terminal domain and the PAZ domain, are involved in small RNA loading and passenger (also called star) strand ejection and 3’ end anchoring of the small RNA, respectively. The two domains of the second lobe, the MID domain and the PIWI domain, are required for the 5’ end binding of the small RNA and the slicing activity of the AGO, respectively. Two additional structured linker domains (L1 and L2) are important for the structural stability of the silencing complex (4). In humans, four AGOs can bind miRNA and siRNA (AGO1, AGO2, AGO3 and AGO4). In *C. elegans*, two AGOs can bind most miRNAs (ALG-1 and ALG-2) (5) and another AGO (ALG-5) can bind a specific subset of miRNAs (6), while multiple other AGOs can bind different types of small RNAs (7). In *Drosophila melanogaster*, Ago1 mainly binds miRNA, while Ago2 mainly binds siRNA because of their respective interactions with specific loading partners (8).

Small RNA loading onto AGOs is an essential step to allow target gene silencing. Several factors are required to achieve an appropriate and specific loading of different types of small RNAs. In *D. melanogaster*, the association between Dicer-2 and R2D2 is required for the specific loading of siRNAs onto Ago2 and to prevent the loading of endogenous siRNA onto Ago1, an AGO generally loaded with miRNAs (9–11). Recent advances in cryo-electron microscopy have allowed a better understanding of the interactions between specific RNA-binding proteins (RBPs) and Dicer, which are involved in small RNA biogenesis and loading. Notably, it was demonstrated how the association between Dcr-2 and Loqs-PD or R2D2 and Dcr-1 and Loqs-PD are necessary for dsRNA processing, siRNA strand selection and pre-miRNA processing, respectively (12–14). The factors regulating small RNA sorting and loading in other animals remain elusive.

The specific loading of different small RNA species into diverse AGOs will specify their function. AGO loaded with miRNAs will generally recruit deadenylation and decapping machinery to inhibit translation and degrade their mRNA targets (15), while AGO loaded with siRNAs will cause their endonucleolytic cleavage (16). The length of the loaded small RNAs, as well as the sequence specificity, can also modulate the miRISC binding outcome, as seen in the slicing efficiency of AGO2 and AGO3, the two slicer-competent human AGOs (17). While some thermodynamic features of the small RNA duplexes (presence of base-pairs mismatches, for example) can influence small RNA sorting into the proper AGOs (18), the factors and mechanisms involved in the differential loading of small RNAs onto AGOs have not been yet completely elucidated. In *C. elegans,* we have previously postulated that intrinsic AGO characteristics, such as structural features or its ability to interact with specific partners, could be required for adequate small RNA sorting among AGOs (19).

Among the factors that may be involved in that process are molecular chaperones, which are potential regulators of AGOs functions (20). Indeed, some of the first interactors of GERp95 (the former name of AGO) identified are chaperones such as Heat-shock protein 70 (HSP70), HSP90 and HSP40 (Hdj-2) (21). Members of the HSP40 family are molecular co-chaperones that interact with HSP70 proteins to modulate their function (22). HSP70 proteins are involved in numerous cellular processes, including proper folding of newly synthesized proteins, preventing protein aggregation and response to different stresses (23). It has been proposed that HSP40 proteins act as molecular guides to dictate the specific function of HSP70 proteins in various cellular contexts (24). Among their numerous functions, molecular chaperones have also been shown to be involved in small RNA pathways. For example, in human cells and *D. melanogaster* extracts, HSP70 (notably the cognate Hsc70-4), HSP90 and an HSP40 co-chaperone are required for small RNA loading (siRNA and miRNA) in an ATP-dependent manner (25–28). Furthermore, single-molecule Förster resonance energy transfer (smFRET) experiments with purified fly Ago2 suggested that HSP70 and HSP40 first bind Ago2 to induce a conformational change and that HSP90 subsequently binds Ago2 to keep it in an open conformation to extend the dwell time of the complex *in vitro* (29). While these biochemical studies suggest the implication of the chaperone machinery in small RNA loading, its involvement in animals has never been assessed.

Here, we identify DNJ-12, a molecular co-chaperone member of the HSP40 family, as a novel key regulator of miRNA loading onto ALG-1 *in vivo*. Despite being a protein with a broad range of binding partners, DNJ-12 interacts with the AGO ALG-1 but not with the other miRNA-specific ALG-2 and other AGOs specific to siRNAs or 21Us. Using the auxin-inducible degron (AID) system, we demonstrate that the tissue-specific depletion of DNJ-12 causes several miRNA-related phenotypes and alters the miRISC ability to repress an mRNA reporter that contains miRNA binding sites in its 3’UTR region, demonstrating that DNJ-12 is required for proper miRNA-mediated gene repression. Furthermore, we show that the loss of DNJ-12 impairs the interaction between ALG-1 and HSP70, a chaperone essential for AGO function *in vitro*. We further demonstrate that the tissue-specific depletion of DNJ-12 leads to the downregulation of specific miRNAs. When depleted ubiquitously, a subset of miRNAs is downregulated and generally less loaded onto ALG-1. We also detect an accumulation of passenger strands, pointing toward an incomplete loading of the miRISC. This study elucidates the role of an HSP40 protein in the specific loading of miRNA in *C. elegans* and outlines the importance of the miRNA loading step in gene regulation by miRNAs in animals.

## MATERIAL AND METHODS

### C. elegans strains and growth conditions

Worm strains were grown on Nematode Growth Media (NGM) 35mm or 150mm plates and fed with OP50 *E. coli* strain. Before performing all experiments, worms were synchronized by treating gravid adult worms with a bleach solution (0.5M KOH, 1.2% [V/V] sodium hypochlorite). The remaining eggs were washed thrice with M9 buffer and kept at 20°C to hatch overnight. Worms were grown at 20°C unless specified otherwise. (Table S1).

### Genome editing

The CRISPR Cas9 system was used to generate multiple worm strains (30). For every strain, approximately 20 worms were injected in one of their gonadal arms with reconstituted Cas9 RNP complex mix (Cas9 (2.5μg/μL), tracrRNA (1μg/μL), guide RNA (0.4μg/μL) and repair template (ssODN) with 35bp homology arms (1.625μM)). dpy-10 was used as an injection efficiency co-marker. All locus edited by the CRISPR Cas9 system were confirmed by Sanger sequencing. The design of guide RNAs was performed using CHOPCHOP V3 (31). Guide RNA and donor template are described in Table S2.

### Protein extraction

Worms were resuspended in 450μL of lysis buffer (100 mM potassium acetate, 30 mM Hepes-KOH pH 7.0, 2 mM magnesium acetate, 1 mM DTT, 0.5% [V/V] Triton X-100, Complete Mini Protease Inhibitor without EDTA (Roche) and 2% [V/V] SUPERase•In) and lysed using a Bullet Blender Storm 24 machine for 5 minutes at maximum speed. Worm lysates were centrifuged for 20 minutes at 17 000g at 4°C, and the cleared supernatant was kept without the lipid phase. Protein concentrations were determined by the Bradford method using the Bio-Rad Protein Assay Dye Reagent Concentrate.

### miRNA complex pull-down

miRNA complex pull-down experiments were done according to Jannot et al (32). 2 mg of worm lysate in each condition were pre-cleared for 1h at room temperature on nutator with M-280 streptavidin Dynabeads coupled with non-specific 2’-*O*-Methyl modified oligonucleotides targeting the luciferase (Luc) sequence. Supernatants were split in two and incubated with either let-7 or Luc 2’-*O*-Methyl modified oligonucleotides for 1h at room temperature. Beads were washed three times with cold lysis buffer. The beads were resuspended in 4x Laemmli loading buffer (1 mM Tris-HCl [pH 6.8], 2% [W/V] SDS, 100mM DTT and 10% [V/V] glycerol) and heated at 99°C for 10 minutes and vortexed 1 minute at top speed using an Eppendorf ThermoMixer C. Boiled samples were loaded on a polyacrylamide gel for Western Blot analysis.

### Immunoprecipitation

10μL of Protein G Dynabeads bound to 2.5μL of anti-HA monoclonal antibody (C29F4, Cell Signaling Technology) or 5μL of anti-ALG-1 polyclonal antibody (homemade) was used per mg of protein lysate. 1 or 2 mg of protein lysate were incubated with the beads for 90 minutes at 4°C (To monitor of FLAG::ALG-2 from DNJ-12::HA immunoprecipitation, we added 10μg RNAse A during the incubation) or 180 minutes for small RNA sequencing experiments. Beads were washed three times with 600μL of cold lysis buffer and resuspended in 4x Laemmli loading buffer. Samples were boiled at 99°C for 10 minutes and vortexed at top speed for 1 minute using an Eppendorf ThermoMixer C. Boiled samples were loaded on a polyacrylamide gel for Western Blot analysis. For immunoprecipitation experiments followed by small RNA sequencing, 2mg of protein was used, and 10% of the immunoprecipitation was loaded on gel for Western Blot validation of the immunoprecipitation efficiency. The rest of the immunoprecipitated ALG-1 was treated with 2μL of Proteinase K for 20 minutes at 50°C followed by TRIzol RNA extraction. All immunoprecipitations were performed either in duplicate (Figure 1B-C) or triplicate (Figure 1D and 5A).

**Figure 1.**
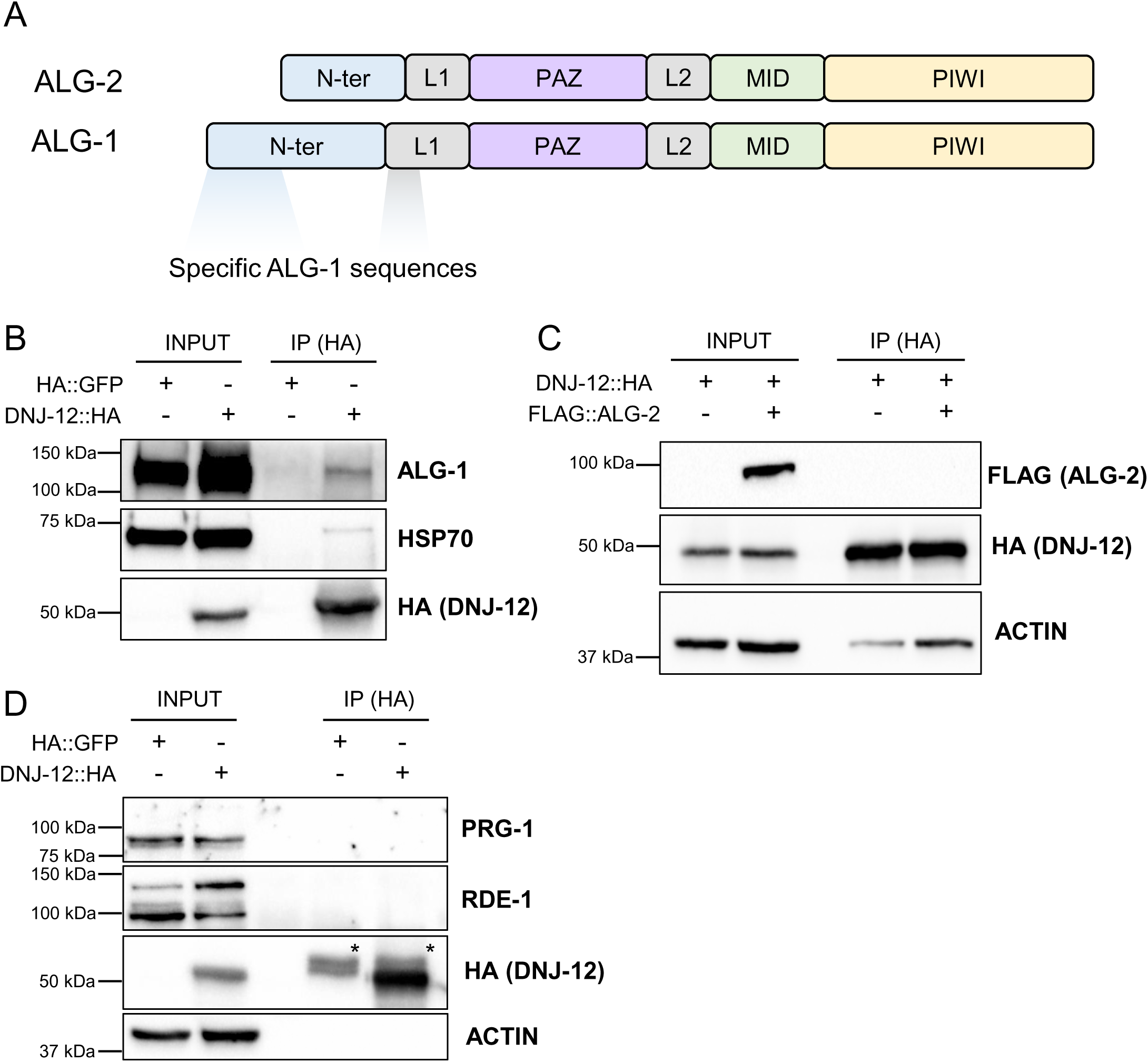
The HSP40 co-chaperone DNJ-12 specifically interacts with the miRNA-specific Argonaute ALG-1. **(A)** Schematic representation of the *C. elegans* miRNA-specific AGOs ALG-1 and ALG-2 domain organization and their distinctive N-terminal domains (see Figure S1 for amino acids alignment). **(B)** Western blot of co-immunoprecipitation of HA::GFP or DNJ-12::HA and ALG-1, HSP-70 and DNJ-12::HA. **(C)** Western blot of co-immunoprecipitation of DNJ-12::HA in the presence or absence of FLAG::ALG-2. Worms were grown at 25°C and collected at the young adult stage. 50ug and 2mg of total lysate were used for input and IP, respectively. **(D)** Western blot of co-immunoprecipitation of HA::GFP or DNJ-12::HA and PRG-1, RDE-1 and ACTIN. * represents Immunoglobulin G.

### Western Blot

Boiled protein samples were analyzed by SDS PAGE on 6% acrylamide stacking gel and 8% acrylamide resolving gel and transferred to 0.45μm nitrocellulose blotting membranes (VWR CA10061-120). Membranes were incubated with primary antibody overnight at 4°C. The primary antibodies used in this paper are anti-ALG-1 (homemade), anti-HSP70 (catalog #SMC-104, StressMarq), anti-HA (catalog # 3724S, NEB), anti-PRG-1 (33), anti-RDE-1(7) and anti-ACTIN (catalog #AB49900, Abcam). Secondary antibodies directed against the primary antibody Sheep anti-mouse IgG HRP (Catalog #515-035-062, Jackson ImmunoResearch Labs), Goat anti-rabbit IgG HRP (Catalog # 111-035-144, Cedarlane) and TrueBlot anti-Rabbit IgG HRP (Catalog #18-8816-31, Rockland) for protein with molecular sizes around 50kDa were used for 1h at room temperature. Membrane imaging was done using Clarity ECL (Catalog #1705061, BioRad), ChemiDoc MP and Image Lab software. In Figure 3A-B, western blots were quantified using the relative quantity tool in Image Lab software. Three replicates were used for the quantification of Figure 3A. The co-immunoprecipitated levels of HSP70 were normalized on input levels of HSP70. Five replicates were used for the quantification of Figure 3B. The levels of ALG-1 and AID::DNJ-12::HA were normalized on the levels of Actin (Table S3).

### Microscopy and imaging

Phenotype images were obtained using a Zeiss AxioCam HRm digital camera mounted on a Zeiss Axio Imager M1 microscope. GFP reporter images were quantified using Zen Software. Worms were mounted on 2% agarose pads and immobilized in a 1% [V/V] levamisole / M9 solution. For IAA treatment efficiency, Box B / ΔN system, and GFP::ALG-1 imaging, a ZEISS LSM 900 confocal microscope was used.

### Quantification of GFP reporter

For the miR-228 reporter, each dot on the bar graphs (Figure 2E-S3B) represents the mean of 4 GFP cells on one worm that were individually quantified using the Zen software. The area used for quantification is outlined by the white dashed lines. For the Box B / ΔN reporter (Figure 3C), each dot represents the quantification of the GFP signal in the terminal bulb of the pharynx of one animal. The area used for quantification is outlined by the white dashed lines (Table S4-S5).

**Figure 2.**
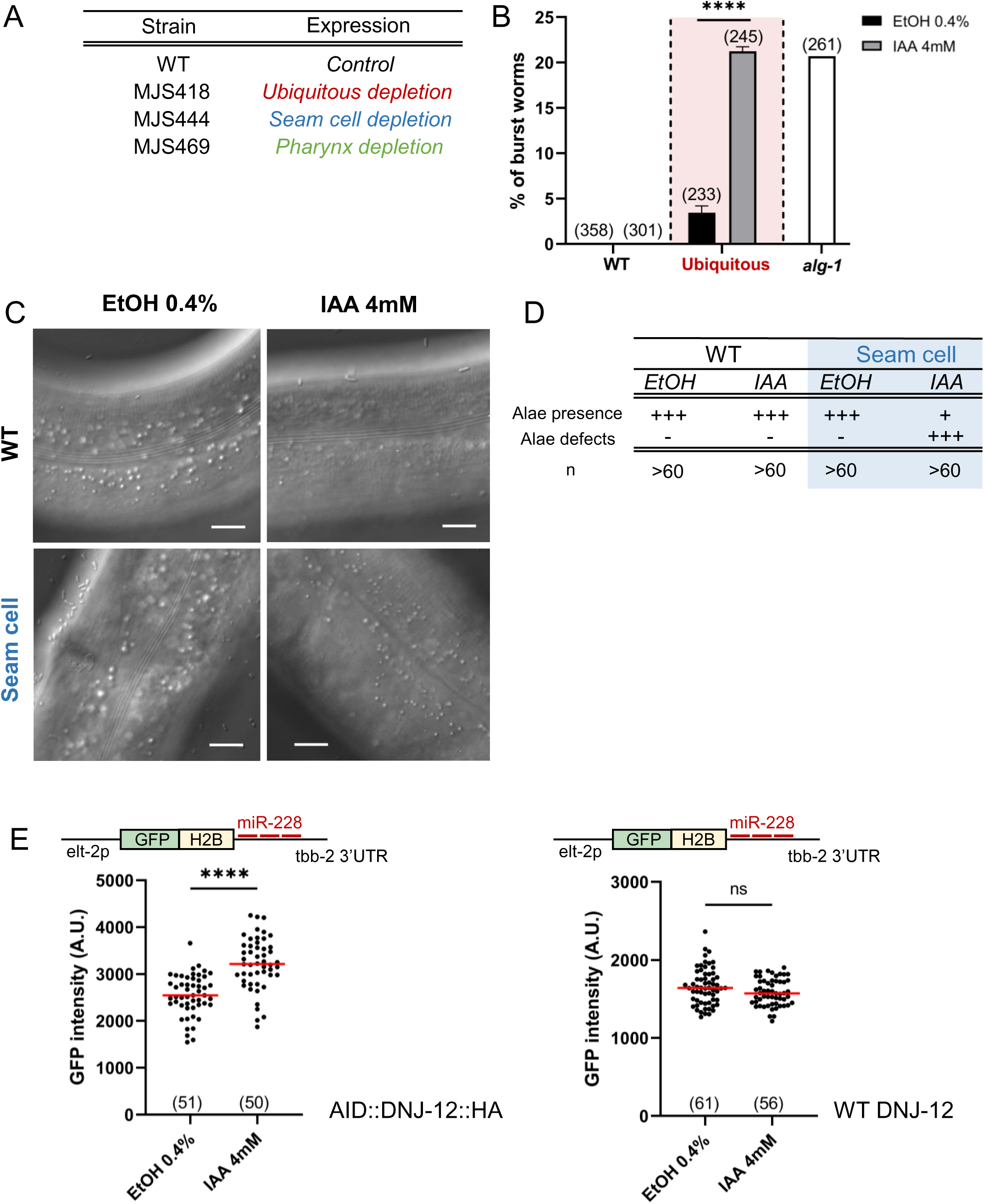
Depletion of DNJ-12 leads to developmental phenotypes and impairs miRNA-dependent repression. **(A**) Table representing the transgenic strains combining AID::DNJ-12::HA and TIR1 protein expressed either ubiquitously (MJS418), in the seam cells (MJS444) or in the pharynx (MJS469). **(B)** Quantification of the vulval bursting phenotype of wild-type (WT) and MJS418 animals grown for their whole post-embryonic development at 20°C on 0.4% vehicle ethanol (EtOH) or 4mM Indole-3-acetic acid (IAA) plates. Three replicates of the EtOH 0.4% treatment or the IAA 4mM treatment were performed. The average percentages of worm bursting are plotted, and the total number of animals scored is in parentheses. The white bar represents *alg-1* (*gk214*) control worms that were grown at 20°C on regular OP50 plates. Error bars represent standard deviation. *p*-value was calculated using Fisher’s exact test. **** represents *p* ≤ 0.0001. **(C)** (Left) DIC microscopy images of the alae of young adults WT and MJS444 worms grown on EtOH 0.4% or IAA 4mM small plates for their whole development at 20°C. The scale bar represents 10μm. (Right) Schematic representation of worms with alae defects. **(D)** Relative quantification of the alae defects phenotypes of wild-type (WT) or MJS444 worms grown on EtOH 0.4% or IAA 4mM plates for their whole post-embryonic development at 20°C. Most MJS444 animals treated with IAA had no observable alae at the young adult stage, while all animals from the other conditions displayed alae at the young adult stage (represented by the + and – symbols). The MJS444 worms treated with 4mM IAA that had detectable alae displayed alae defect (represented by +++), while worms from the other conditions had no alae defect. **(E)** (Left) Quantification of the GFP signal in arbitrary units of three biological replicates of auxin treatments (MJS482 worms). In each biological replicate, each dot represents the mean of 4 GFP cells on one worm. The horizontal bar represents the mean. *p*-value was calculated using a two-tailed unpaired t-test. **** represents *p* ≤ 0.0001. (Right) Quantification of the GFP signal in arbitrary units of three biological replicates of auxin treatments (MJS488 worms). In each biological replicate, each dot represents the mean of 4 GFP cells on one worm. The horizontal bar represents the mean. *p-*value was calculated using a two-tailed unpaired t-test. ns represents *p* > 0.05.

### RT-qPCR

Synchronized worms were collected with M9 and left for 30 minutes with agitation at room temperature. Worm pellets were washed 3 times with M9. 4 volumes of TRIzol were added to the samples, flash frozen and vortexed 1 minute at maximum speed using an Eppendorf ThermoMixer C. RNA extraction was performed with chloroform and precipitated with ethanol 100% [V/V] overnight at -20°C. The RNA pellet was washed with ethanol 75% [V/V] and solubilized in 20μL of RNase-free water. RNA quantification was performed with a NanoDrop One. For miRNA quantification, reverse transcription was performed using 10ng of RNA with Applied Biosystems Small RNA Assay protocol (16°C for 30 minutes, 42°C for 30 minutes and 85°C for 5 minutes). Taqman RT-qPCR was performed using 1.33μL of cDNA on a QuantStudio 3 machine with regular settings (enzyme activation at 95°C for 10 minutes and 40 cycles at 95°C for 15 seconds and 60°C for 60 seconds). sn2841 was used as an endogenous control to normalize miRNA levels. For mRNA quantification, reverse transcription was performed using 400ng of RNA with Multiscribe Reverse Transcriptase Kit (Applied Biosystems) (25°C for 10 minutes, 37°C for 120 minutes and 85°C for 5 minutes) followed by RNase H treatment (1uL per RT) at 37°C for 30 minutes. SYBR Green RT-qPCR was performed using 2uL of cDNA with the SsoAdvanced Universal SYBR Green Supermix (Bio-Rad) on a QuantStudio 3 machine with regular settings (95°C for 10 minutes and 40 cycles at 95°C for 15 seconds and 60°C for 60 seconds). Tba-1 was used as an endogenous control to normalize mRNA levels. All RT-qPCR experiments were performed in triplicate. Analyses were performed using the ΔΔCt method.

### Sequence alignment

Alignment of the N-terminal region of both isoforms of ALG-1 and ALG-2 (Figure S1), as well as the alignment of *Arabidopsis thaliana* AGO10, *Caenorhabditis elegans* ALG-1 and human AGO2, and human AGO1-4 (Figure S8A-S8B) were performed using Clustal Omega on default settings (34).

### Small RNA sequencing

To clone the small RNA libraries, we used the NEBNext® Multiplex Small RNA Library Prep Kit for Illumina (E7560S) with RNA extracted directly from total worm lysates or after ALG-1 IP (Supplementary Tables S6-S6-S7-S9-S10-S11-S12). NovaSeq 6000 S1 (v1.5) reads were mapped to the genome and cDNA using custom PERL (5.10.1) scripts and Bowtie 0.12.7 (35). Databases used include the *C. elegans* genome (WormBase release WS215), Repbase 15.10 (36), and miRbase 16 (37). The samples were normalized to total gene expression (geex) reads excluding structural RNA (struc) and transformed in RPM (read per million). The mean log_2_ value of three biological replicates for every condition was used to generate the graphs. The principal component analysis confirmed that replicates are similar (Figure S5A). The sequencing data are available on Gene Expression Omnibus (GEO) using the accession number GSE241034.

### Statistical analysis

In Figures 2E, 3C, 4A, S3B and S3D, two-tailed unpaired t-tests were performed using GraphPad Prism 9 software. In Figure 2B, Fisher’s exact test was performed using GraphPad 9. Results are plotted as follows: ns represents *p* > 0.05, * represents *p* ≤ 0.05, ** represents *p* ≤ 0.01, *** represents *p* ≤ 0.001 and **** represents *p* ≤ 0.0001.

### Indole-3-acetic acid treatment (IAA)

Before every IAA treatment, worms were synchronized using the bleaching protocol previously described. The IAA treatment was done using NGM plates supplemented with IAA 4mM or EtOH 0.4% [V/V] as a control. IAA or EtOH were added while the NGM was cooling (between 40°C and 45°C). Concentrated OP50 bacteria were added to the plate the next day, and the plates were stored at 4°C until they were used. The IAA plates were kept for a maximum of 2 weeks.

For the worms grown at 20°C, we noticed a minimal amount of bursting animals (0.4%) in the strain expressing AID::DNJ-12::HA and TIR1 under *eft-3* promoter (Figure S3C). Since the individual mutant strain did not display any bursting phenotype, we considered that AID::DNJ-12::HA is functional at this temperature (Figure S3C). However, we noticed that the worms grown at 25°C displayed a modest level of bursting phenotype (between 0% and 5%), indicating that AID::DNJ-12::HA function might be slightly impaired at 25°C (Figure S3C). Therefore, we conducted all IAA depletion experiments at 20°C to avoid possible non-specific effects.

Synchronized larvae were put on seeded IAA plates and grown at 20°C for either 30 minutes, 60 minutes, 180 minutes or until they reached the L4 or young adult stage, depending on the experiment. The efficiency of depletion was either monitored by western blot in ubiquitous depletions or by confocal microscopy in tissue-specific depletions (indirect measurement of TIR1 activity using the internal BFP::AID::NLS tagged control localized after the F2A self-cleaving peptide (38)).

## RESULTS

### The HSP40 co-chaperone DNJ-12 specifically interacts with miRNA-specific AGO ALG-1

In *C. elegans*, the two principal miRNA-associated AGOs, ALG-1 and ALG-2, share up to 88% of sequence identity depending on the isoforms (39). The main differences between both AGOs reside in the more extended N-terminal domain and L1 region of ALG-1 (Figure 1A-S1). Indeed, ALG-1 and ALG-2 are almost identical from the PAZ to the PIWI domain (93% identity). Since both AGOs have overlapping yet distinct functions (39, 40), we hypothesized that the unique N-terminus of ALG-1 interacts with specific protein(s) that can confer its exclusive functions. To identify these specific interactors, we previously performed a yeast two-hybrid screen using the first 132 amino acids of ALG-1 as bait (41) (Figure S1). Among the interacting partners, we identified DNJ-12, a constitutively express HSP40 co-chaperone (42). To validate that the interaction between ALG-1 and DNJ-12 occurs in animals, we generated a *C. elegans* strain carrying C-terminal HA-tagged alleles of *dnj-12* at the endogenous locus using the CRISPR Cas9 genome editing method to overcome the absence of specific antibodies. This tagged strain does not display any specific phenotypes and is superficially wild-type. We performed a co-immunoprecipitation of DNJ-12::HA or the negative control HA::GFP and detected an interaction between DNJ-12 and ALG-1, confirming that ALG-1 interacts with DNJ-12 *in vivo*. (Figure 1B). We also detected an interaction between DNJ-12 and HSP70, a canonical interactor of DNJ-12 (Figure 1B). Since HSP40 can transiently bind a large subset of client proteins due to their co-chaperone activity (43), we sought to test whether DNJ-12 is also binding to other AGOs. First, we co-immunoprecipitated DNJ-12::HA and FLAG::ALG-2 (no antibodies are available for the detection of ALG-2) in presence of RNAse to eliminate potential indirect interaction caused by mRNA targets bound by both ALG-1 and ALG-2 miRISCs, and we were not able to detect any association between both proteins (Figure 1C), which is in line with the fact that ALG-2 lacks the extended N-terminal region specific to ALG-1 (Figure S1). Furthermore, we also monitored the interaction between DNJ-12::HA and RDE-1 or PRG-1, two AGOs involved in RNAi and 21U-mediated silencing, respectively (44–46), by immunoprecipitation and we were unable to detect any interaction (Figure 1D). Taken together, our data indicate that DNJ-12 is a novel interactor of ALG-1 and does not interact with ALG-2, RDE-1 and PRG-1.

### Depletion of DNJ-12 leads to developmental phenotypes and alters miRNA function

Since miRNAs are important regulators of several developmental cues throughout the nematode life, we wanted to investigate the potential contribution of DNJ-12 in these regulatory events. Some of the most established developmental processes regulated by miRNAs are the proliferation and differentiation of seam cells. The seam cells are specialized hypodermal cells that divide through post-embryonic development. At the L4 to adult transition, the 16 seam cells fuse to generate a syncytium and give rise to alae, cuticular ridges spanning on two sides of the animals (47, 48). Specific miRNAs tightly regulate the terminal fusion of seam cells along with the generation of alae. Notably, the alteration of the let-7 miRNA family (let-7, miR-48, miR-84, miR-241) and many components of the miRNA pathway, such as *alg-1*, leads to an increased number of seam cells and defects in the alae formations (49–51). Also, miRNAs play a central role in the nematode vulval development. In this case, misregulation of *lin-41* mRNA by let-7 leads to vulval bursting (49, 52).

To test the contribution of *dnj-12* in these miRNA-dependent phenotypes, we first planned to use a loss-of-function mutant strain. However, the complete knockout of *dnj-12* is lethal (53), likely due to its involvement in several important processes. To circumvent that problem, we chose to use the newly characterized degron system recently optimized in *C. elegans*, allowing a rapid and acute depletion of any protein of interest (38). This system includes *Arabidopsis thaliana* TIR1 protein, which is an F-box protein that can interact with endogenous proteins to generate a functional ubiquitin ligase complex that is activated by auxin (IAA) treatment, as well as an endogenously tagged version of DNJ-12 (AID::DNJ-12::HA) that we generated using the CRISPR Cas9 system. Furthermore, the strains expressing TIR1 under different promoters also contain an internal control, mTagBFP2::AID (blue fluorescent protein), that allows the monitoring of the activation of the degradation system after IAA treatment (38). To degrade AID::DNJ-12::HA in the whole worm, we generated a *C. elegans* strain (MJS418) carrying AID::DNJ-12::HA and expressing TIR1 under the promoter *eft-3*, a ubiquitously expressed translation elongation factor (38). Additionally, we also generated two other strains where DNJ-12 can be degraded, specifically in the seam cells (MJS444) and in the pharynx (MJS469), to study the involvement of DNJ-12 in these specific tissues (Figure 2A). We chose to deplete DNJ-12 in the pharynx because it is one of the tissues where ALG-1 is strongly expressed (39). We confirmed that the TIR1 enzyme was active upon treatment with 4mM IAA by observing the loss of the BFP signal in these tissues (Figure S2A-S2B). Furthermore, we also validated the efficiency of AID::DNJ-12::HA depletion after IAA treatment by western blot and confirmed that AID::DNJ-12::HA was almost entirely degraded after 180 minutes (Figure S2C).

First, we wanted to know if the ubiquitous loss of DNJ-12 affected the function of the let-7 miRNA by looking at the vulval bursting (Figure S2D), a phenotype caused by the loss of let-7 function in *C. elegans* (49, 54). Accordingly, we exposed synchronized MJS418 animals to either control treatment (0.4% ethanol) or 4mM IAA for their entire post-embryonic development. The treated worms showed about 20% of bursting, comparable to the level of *alg-1(gk214)* null allele, while the control worms displayed less than 5% of this phenotype (Figure 2B). This result suggests that the loss of DNJ-12 leads to abnormal gene regulation by the let-7 miRNA. Next, we decided to study the implication of DNJ-12 on seam cells development by using the *C. elegans* strain in which DNJ-12 is only depleted in these cells. Interestingly, we observed that the IAA treatment almost completely prevents the alae formation, and for the few ones that still formed alae, severe gaps and breaks were observed (Figure 2C-2D; no normal alae were detected in treated animals). These results indicate that DNJ-12 is important in regulating developmental processes regulated by miRNAs.

To investigate if DNJ-12 was required for the function of other miRNA families, we used a GFP reporter under the control of miR-228 (55), depleted DNJ-12 ubiquitously and monitored GFP levels in intestinal cells. We observed that the depletion of DNJ-12 caused a significant increase in the GFP signal compared to the control treatment, indicating that DNJ-12 is required for proper miR-228-mediated target repression (Figure 2E-S3A). To validate that the IAA treatment and the activation of TIR1 under the *eft-3* promoter were not affecting the reporter expression on their own, we monitored the GFP signal in a strain without the AID-tagged DNJ-12 (MJS488) and did not detect any significant differences in GFP signal from worms treated with EtOH or IAA, suggesting that the dysregulation of the miR-228 reporter is caused by the DNJ-12 depletion (Figure 2E-S3A). Using a strain containing mutated miR-228 binding sites, we confirmed that most GFP derepression depends on miR-228 binding (Figure S3B). We also monitored the level of four validated endogenous miRNA targets (49, 54, 56–58), and we were able to detect increased levels of mRNAs upon DNJ-12 depletion (Figure S3D), suggesting that the loss of DNJ-12 also impairs the miRNA-dependent repression of endogenous targets. Altogether, these data support that the co-chaperone DNJ-12 participates in the miRNA-mediated gene regulation in *C. elegans*.

### DNJ-12 contributes to ALG-1 association with the molecular chaperone HSP70 but does not affect its gene silencing capacity

To uncover the underlying molecular mechanisms involved in the regulation of ALG-1 by DNJ-12, we further characterize the interaction between DNJ-12 and the miRISC. To first determine if DNJ-12 interacts with the repressive miRISC (miRISC loaded with a miRNA), we performed a miRISC pull-down using a probe complementary to let-7 miRNA sequence (32). We detected the presence of ALG-1, but not DNJ-12, suggesting that DNJ-12 does not interact with the active miRISC (Figure S4A).

Since HSP40 can act as molecular guides to modulate HSP70 interactions with client proteins, we wondered if the loss of DNJ-12 could impact the interaction between ALG-1 and the molecular chaperone HSP70. To answer this question, we performed immunoprecipitation of ALG-1 after AID::DNJ-12::HA ubiquitous depletion. While we detect an association between HSP70 and ALG-1 in normal conditions (Figure 3A), the depletion of DNJ-12 greatly impairs their interaction, implying that DNJ-12 is important for the interaction between ALG-1 and HSP70, which is in accordance with its co-chaperone function (Figure 3A).

**Figure 3.**
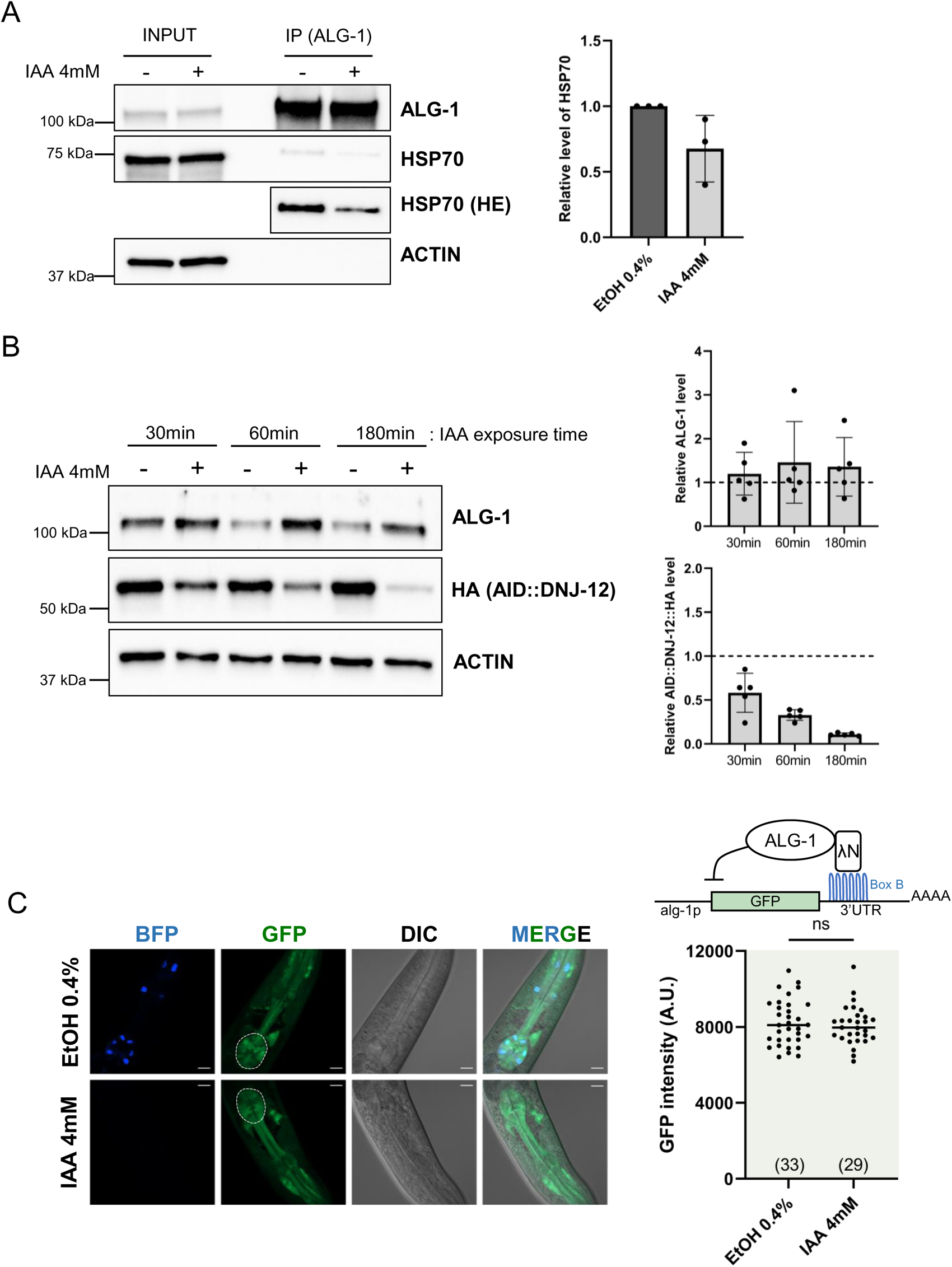
DNJ-12 modulates ALG-1 interaction with HSP70 without affecting its gene silencing capacity. **(A)** (Left) Western blot of co-immunoprecipitation between ALG-1 and HSP70. Worms were grown at 20°C on either EtOH 0.4% plate or IAA 4mM plates and collected at the young adult stage. HE means higher exposure. 50ug and 2mg of total lysate were used for input and IP, respectively. (Right) Quantification of the levels of HSP70 interaction with ALG-1 from three replicates of ALG-1 immunoprecipitation. The co-immunoprecipitated levels of HSP70 were normalized on input levels of HSP70. Error bars represent standard deviation. **(B)** (Top) Western blot of worms boiled in Laemmli buffer (MJS418) after 30 minutes, 60 minutes or 180 minutes of IAA 4mM treatment and detection of ALG-1, AID::DNJ-12::HA and ACTIN. Worms were grown for 55h at 20°C, then shifted on IAA 4mM small plates for the indicated time. (Bottom) Quantification of the levels of AID::DNJ-12::HA and ALG-1 from five replicates after three different IAA treatment times: 30 minutes, 60 minutes or 180 minutes. AID::DNJ-12::HA and ALG-1 levels were normalized on ACTIN levels. Error bars represent standard deviation. **(C)** (Left) Representative confocal images (BFP, GFP, DIC and merge) of L4 MJS475 worms (expressing GFP with Box B sequences, λN::alg-1, AID::DNJ-12::HA and pharyngeal TIR1) grown on either EtOH 0.4% or IAA 4mM plates for their whole development at 20°C. The scale bars represent 10μm. (Right) Quantification of the GFP signal in arbitrary units of three biological replicates of auxin treatment experiments. Each dot represents the quantification of the GFP signal in the terminal bulb of the pharynx of one animal. The horizontal bar represents the mean. *p*-value was calculated using a two-tailed unpaired t-test. ns represents *p* > 0.05.

Since the HSP70/HSP40 machinery can regulate the folding state of several proteins, we wanted to investigate if DNJ-12 could regulate ALG-1 stability. To do so, we depleted AID::DNJ-12::HA ubiquitously for 30, 60 or 180 minutes to study the impact on ALG-1 stability. AID::DNJ-12::HA is already strongly degraded after 30 minutes of IAA treatment and after 180 minutes, there is almost no AID::DNJ-12::HA left in the worms (Figure 3B). We noticed that although we could detect an increase in ALG-1 levels, the results of these experiments were highly variable, ranging from an important effect on ALG-1 stability to no effect at all in independent experiments (Figure 3B).

To further investigate the impact of depleting DNJ-12 on ALG-1, we wanted to test if ALG-1 can effectively repress a target without the need of a miRNA. To do so, we use an artificial tethering system that we previously generated where ALG-1 is fused to a λN peptide and tethered to a GFP reporter containing Box B sequences (Figure 3C) (59). The repression of this reporter by ALG-1 is independent of the interaction between mRNA and miRNA, and thus independent of miRISC loading state. We also previously validated that this GFP reporter is derepressed in the absence of ALG-1 tethering, confirming that the expression of the reporter is dependent on ALG-1 repression (59). To use our tethering system combined with the IAA-depletion of DNJ-12, we generated a new worm strain (MJS475) containing Box B GFP reporter, λN::ALG-1 and AID::DNJ-12::HA with a TIR1 protein specifically expressed in the pharynx (MJS469). We compared the GFP expression of worms grown on either control or IAA plates and did not notice any difference in the expression of the reporter, suggesting that the miRISC is still able to perform mRNA silencing in the absence of DNJ-12 when tethered to a GFP reporter (Figure 3C).

Collectively, these data suggest that the co-chaperone DNJ-12 only interacts with a small fraction of the ALG-1 complexes and that it does not interact with the miRNA-bound miRISC. Also, DNJ-12 mediates ALG-1 interaction with the chaperone HSP70 and potentially regulates ALG-1 stability. ALG-1 can still repress a GFP reporter when artificially tethered to it, suggesting that ALG-1 is still functional and able to recruit key partners in the absence of DNJ-12. Hence, the regulation of ALG-1 function by DNJ-12 does not occur at the level of recruitment of the silencing machinery but rather at a step preceding miRISC binding to its target, pointing toward a possible role in the miRISC loading by miRNAs.

### Depletion of DNJ-12 leads to a decrease in microRNA levels and their loading onto ALG-1 Argonaute

Because DNJ-12 depletion impairs miRISC function and the interaction between DNJ-12 and ALG-1 does not occur at the repressive miRISC stage, we hypothesized that DNJ-12 could contribute to miRNA loading onto ALG-1 AGO *in vivo*. This hypothesis is in line with *in vitro* studies suggesting that the HSP70 machinery and the HSP90 machinery, enhance the miRISC loading through AGO remodelling (25–28). To test the role of DNJ-12 on ALG-1 miRNA loading, we first examined the level of several miRNAs after DNJ-12 depletion by focussing our initial characterization on miRNAs that were predicted to be expressed specifically in our depleted tissues. Hence, we monitored the levels of let-7, miR-244 and miR-255, expressed ubiquitously, in the seam cells and the pharynx, respectively (60) (Figure 4A). After the ubiquitous depletion of DNJ-12, we were able to detect a significant decrease in let-7 and miR-255 levels (*p*-value=0.021 and 0.022, respectively) and a slight decrease in miR-244 level (*p*-value=0.063) (Figure 4A). In the seam cells, we were able to detect a significant recduction of miR-244 (*p*-value=0.029) and in the pharynx, of miR-255 (*p*-value=0.0080) (Figure 4A). When we analysed the global levels of miRNAs upon tissue-specific DNJ-12 depletions by performing small RNA sequencing from three biological replicates, we observed that only one miRNA (0.8% of all miRNAs) was downregulated (Δlog_2_ RPM of -0.5) in the seam cells, 13 miRNAs were downregulated in the pharynx (10.4% of all miRNAs) and 28 miRNAs were downregulated ubiquitously (22.4% of all miRNAs) (Figure 4B-S5B). Among the ubiquitously downregulated miRNAs, nearly all (24/28) were also shown to be downregulated significantly in *alg-1 (gk214)* mutant (61). On the other hand, 9 miRNAs (7.2% of all miRNAs) were upregulated in the seam cells, 8 were upregulated in the pharynx (6.4% of all miRNAs) and 2 were upregulated ubiquitously (1.6% of all miRNAs).

**Figure 4.**
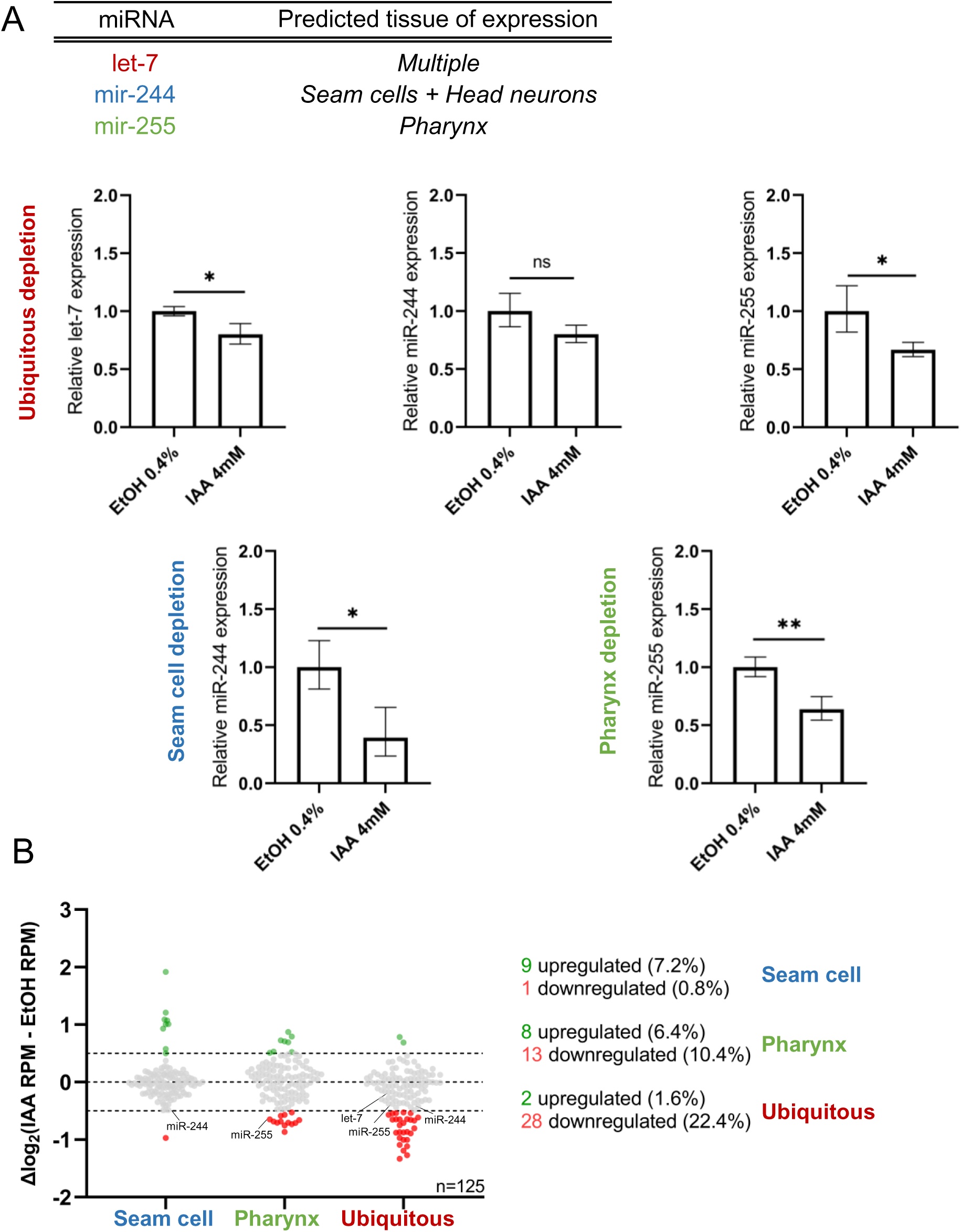
Tissue-specific depletion of DNJ-12 leads to a decrease in miRNA levels. **(A)** (Top) Table presenting multiple miRNAs and their predicted tissues of expression. (Bottom) Quantification of miRNA (let-7, miR-244 or miR-255) expression measured by RT-qPCR. RNA was purified from young adult worms grown on EtOH 0.4% or IAA 4mM plates for their whole development at 20°C. The small nucleolar RNA sn2841 was used as an endogenous control to obtain ΔΔCT values. The error bars indicate the 95% confidence interval. *p*-values were calculated using a two-tailed unpaired t-test. ns represents *p* > 0.05, * represents *p* ≤ 0.05 and ** represents *p* ≤ 0.01. **(B)** miRNA sequencing data plotted according to IAA-mediated DNJ-12 depletion in the seam cells, the pharynx or ubiquitously. The raw reads of three biological replicates were individually normalized on the total gene expression reads (geex) minus the structural reads (struct) and converted to RPM. The average log_2_ RPM values of vehicle ethanol (EtOH)-treated worms were subtracted from the average log_2_ RPM values of IAA-treated worms to generate the Δlog_2_ RPM values that were plotted. Each dot represents one miRNA. Only miRNAs that have a normalized RPM above 1 in every sample were kept for the analysis. Only guide strands are presented in this graph.

To confirm that DNJ-12 is not involved in the regulation of other AGOs, we monitored the level of different small RNAs: 21U-RNAs loaded by PRG-1 and 26G-RNAs loaded by ALG-3/ALG-4 (ERGO-1 26Gs cannot be assessed as they are hardly expressed in young adult animals (62)). Our sequencing data demonstrates that neither 21U-RNAs nor 26G-RNAs were impacted by DNJ-12 depletion (Figure S6A). Given that alterations in these AGOs often lead to a decrease in the level of their bound small RNAs (33, 63–65), our sequencing data suggest that these pathways remain unaffected following DNJ-12 depletion. Collectively, these results suggest that targeted depletion of DNJ-12 may destabilize certain miRNAs expressed in specific tissues while leaving other small RNA pathways unaltered.

Knowing that AGO loading by small RNAs stabilizes both AGO and small RNAs (66, 67) and that preventing this molecular step causes destabilization and degradation of miRNAs, as observed in the *alg-1* null animals where several miRNA levels are strongly decreased (50, 61, 68), we investigated if the loading of some miRNAs was affected by the depletion of DNJ-12. To answer this question, we depleted DNJ-12 ubiquitously and performed ALG-1 immunoprecipitations before small RNA sequencing only to detect AGO-loaded miRNAs. Interestingly, 39 miRNAs (29.6%) were less loaded (Δlog_2_ of -0.5), suggesting that DNJ-12 is involved in the loading of a subset of miRNAs (Figure 5A). Next, to further extend our understanding of the relationship between miRNAs variations and their loading states, we overlapped the results from our total and immunoprecipitated small RNA sequencing experiments. Among the 39 miRNAs that were downregulated in the immunoprecipitation condition, 24 (61.5%) were also downregulated in the total miRNAs condition (Figure 5B-S6B). Conversely, among the 12 miRNAs that were upregulated in the immunoprecipitation condition, only 1 (8.3%) was also upregulated in the total miRNAs condition (Figure 5B). Moreover, we compared the expression of several miRNA star strands with their corresponding guide strand to better understand the miRNA duplex dynamic. Globally, we observe a trend between the levels of decreased star strand and their corresponding guide strand in both total and immunoprecipitated conditions (Figure 5C-5D). On the other hand, there is no such trend between the level of upregulated star strand and their corresponding guide strand in both conditions (Figure 5C-5D). The reduction of many miRNA duplexes observed in the absence of DNJ-12 strongly suggests its implication in duplex loading onto ALG-1. The increased loading of a subset of miRNA star strands onto ALG-1 infers that the loss of DNJ-12 might also impair the star strand ejection from the AGO.

**Figure 5.**
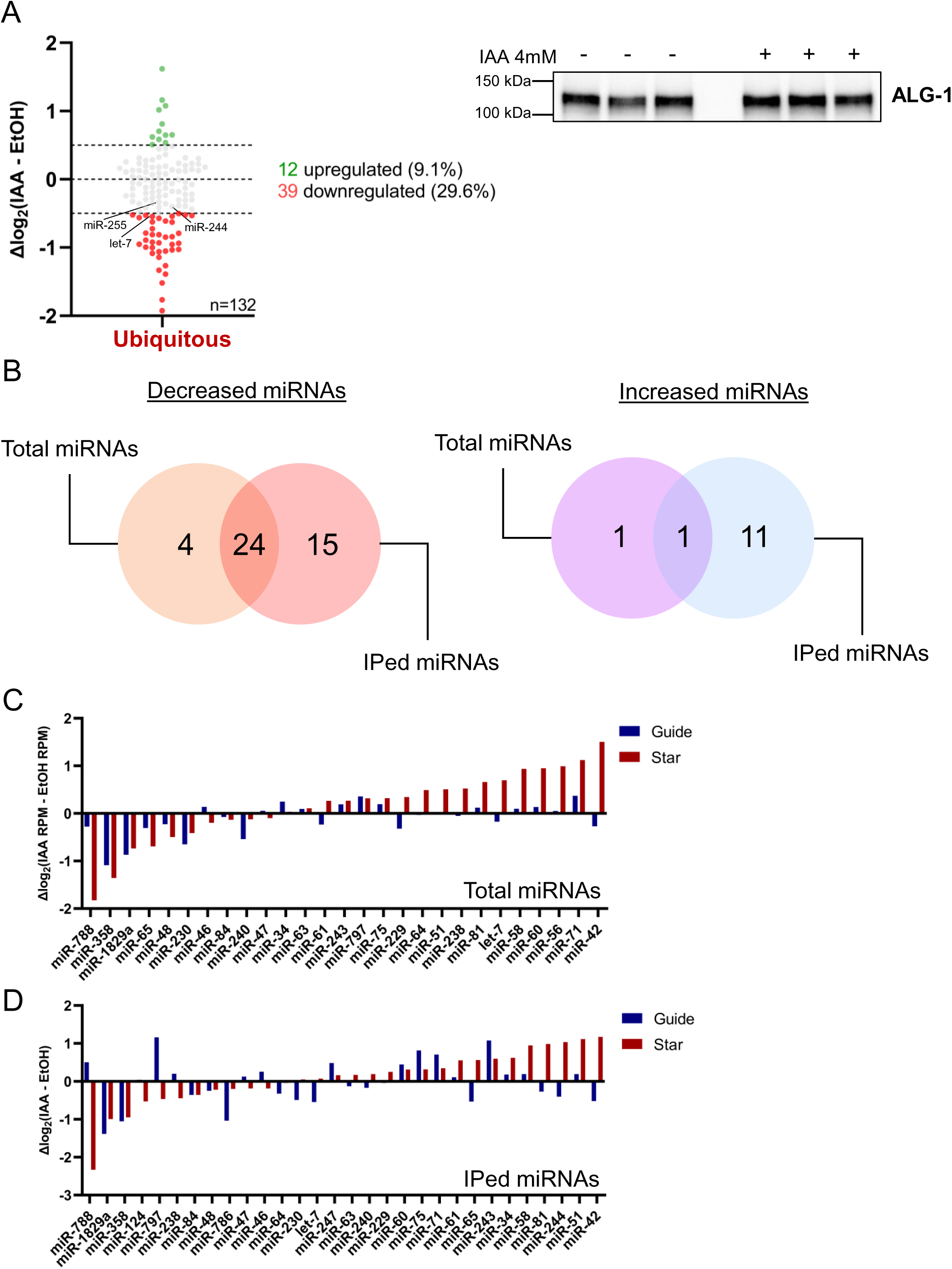
DNJ-12 depletion impairs miRNA loading and miRNA* levels. **(A)** (Left) miRNA sequencing data after DNJ-12 ubiquitous depletion and ALG-1 immunoprecipitation. The raw reads of three biological replicates were individually normalized on the level of ALG-1 protein immunoprecipitated. The average log_2_ values of EtOH-treated worms were subtracted from the average log_2_ values of IAA-treated worms to generate the Δlog2 values that were plotted. (Right) Western blot showing the results of three biological replicates of ALG-1 immunoprecipitation. 10% of the IP (2mg) was loaded for western blot validation, and 90% of the IP was used for small RNA sequencing library preparation. **(B)** (Left) Venn diagram showing the overlap between total miRNAs and loaded miRNAs downregulated after ubiquitous depletion of DNJ-12 by IAA treatment. 28 miRNAs are downregulated (-0.5 log_2_ RPM) in the total extract and 39 miRNAs are downregulated after ALG-1 immunoprecipitation. 24 microRNAs are downregulated in both conditions. (Right) Venn diagram showing the overlap between total and loaded miRNAs upregulated after ubiquitous depletion of DNJ-12 by IAA treatment. 2 microRNAs are upregulated (+0.5 log_2_ RPM) in the total extract and 12 miRNAs are upregulated after ALG-1 immunoprecipitation. 1 miRNA is upregulated in both conditions. **(C)** Bar graph representing the total levels of microRNA guide strands and their corresponding star strand after ubiquitous depletion of DNJ-12 by IAA and small RNA sequencing. Each bar represents the average log_2_ RPM of 3 biological replicates of total microRNAs from EtOH-treated worms subtracted from the average log_2_ RPM of 3 biological replicates of total miRNAs from IAA-treated worms. Only the detectable star strands and their complementary guide strands were plotted. **(D)** Bar graph representing the loaded levels of miRNA guide strands and their corresponding star strands after ubiquitous depletion of DNJ-12 by IAA and small RNA sequencing. Each bar represents the average log_2_ of 3 biological replicates of loaded microRNAs from EtOH-treated worms subtracted from the average log_2_ of 3 biological replicates of loaded microRNAs from IAA-treated worms. Only the detectable star strands and their complementary guide strands were plotted.

Taken together, these results suggest that DNJ-12 is important for loading specific miRNA duplexes onto ALG-1.

## DISCUSSION

Here, we identified the co-chaperone DNJ-12 as a new interactor of the miRNA-specific AGO ALG-1 in *C. elegans*. We also demonstrated that the depletion of DNJ-12 alters the capacity of miRNAs to regulate mRNA targets in animals properly. Furthermore, we found that the interaction between the molecular chaperone HSP70 and ALG-1 is strongly diminished when DNJ-12 is depleted, suggesting that this co-chaperone participates in ALG-1 interactions with binding partners. Functionally, we demonstrated that DNJ-12 regulates miRNA loading onto ALG-1. When we depleted DNJ-12, we detected a decrease in the loading of a subset of miRNA duplexes and, consequently, their downregulation systemically. We also detect an increase in the loading of some passenger strands without any effect on their corresponding guide strand loading, pointing toward a passenger strand ejection defect. These results suggest a model where ALG-1 might be unable to undergo conformational changes to accommodate the loading of specific duplexes due to its loss of interaction with the HSP70 machinery (Figure 6).

**Figure 6.**
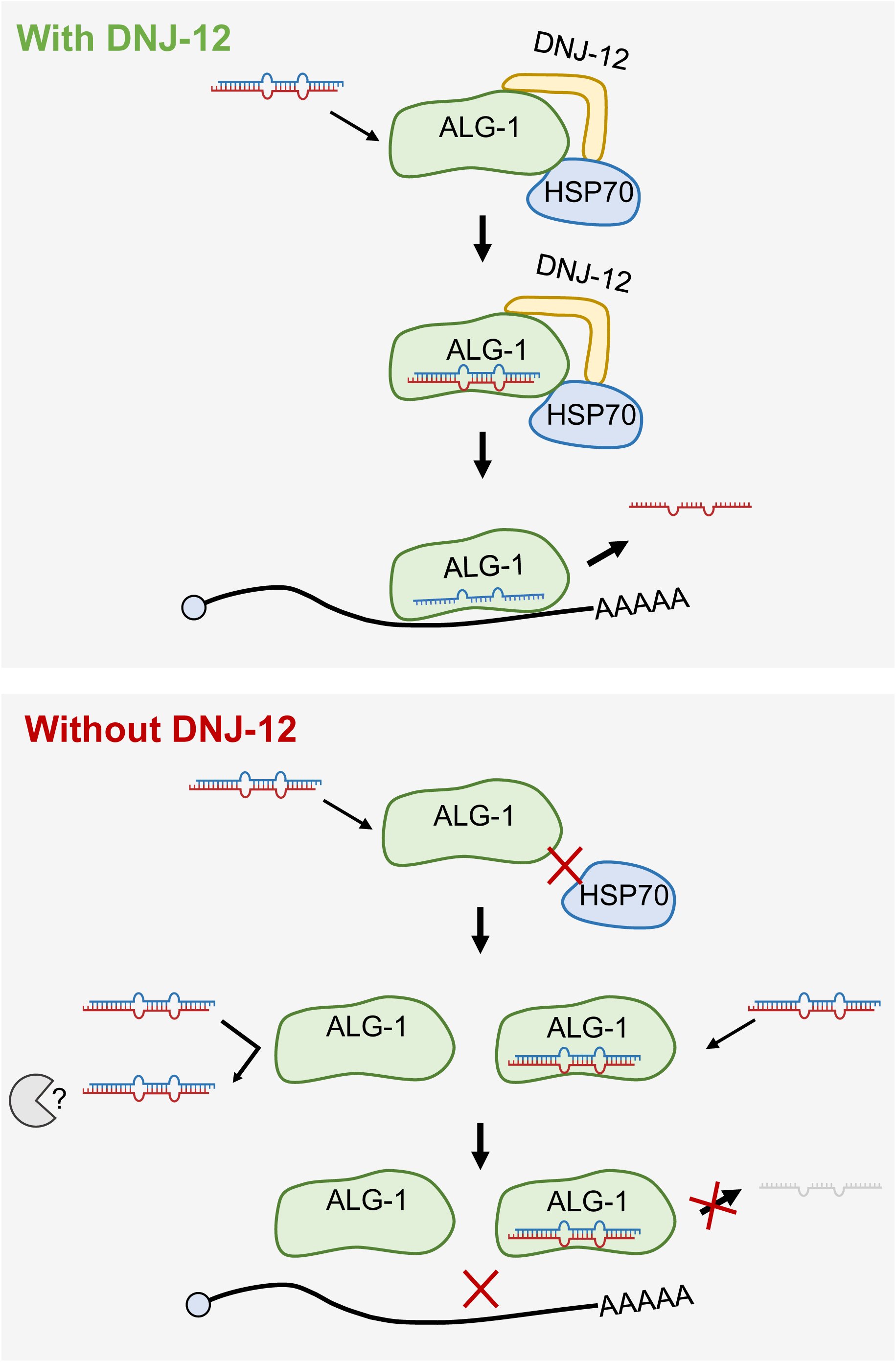
DNJ-12 regulates ALG-1 loading by a subset of miRNAs. **(A**) In normal conditions, DNJ-12 interacts with ALG-1 and promotes the interaction between HSP70 and ALG-1. The miRNA duplexes can then be loaded on ALG-1 to form the miRISC. The miRISC can bind to mRNA targets and regulate their expression. In conditions without DNJ-12, the interaction between HSP70 and ALG-1 is diminished. A subset of miRNA duplexes is cannot be loaded on ALG-1 or to eject the passenger strand, preventing the adequate regulation of mRNA targets.

How can DNJ-12 molecularly affect ALG-1 AGO? When we monitored ALG-1 levels upon DNJ-12 depletion, we observed variable effects on ALG-1 accumulation (Figure 3B). One possibility to explain this phenomenon is that the absence of DNJ-12 increases the amount of empty ALG-1 in animals. It has been reported that unloaded AGOs (or AGOs that cannot eject passenger strands) are degraded by the proteasome and selective autophagy (69, 70). While the mechanism underlying the degradation of empty (or Apo) AGO in *C. elegans* is still unknown, we can envision that DNJ-12 is contributing to ALG-1 conformational changes that will have an incidence on the amount of different AGO states (e.g. Apo, pri-miRISC, pre-miRISC and miRISC (20)), and thus its homeostasis. It is also possible that the interaction with DNJ-12 could regulate ALG-1 subcellular localization. HSP40 can be subjected to a post-transcriptional modification named farnesylation. This type of modification requires a CAAX (C=Cysteine, A=Aliphatic amino acid and X= Any amino acid) motif at the C-terminus of its target proteins. Interestingly, DNJ-12 possesses a similar motif (CQQS) and its orthologs also possess similar terminal motifs: CQTS in DNAJA1 (*H. sapiens*), CTSS in Droj2 (*D. melanogaster*), CASQ in YDJ1 (*S. cerevisiae*) and CAQQ in J2 and J3 (*A. thaliana*). Moreover, DNAJA1, YDJ1 and J2/J3 have been shown to be farnesylated experimentally (71–73), suggesting that it could also be the case for DNJ-12. Remarkably, it was demonstrated that J2 and J3 were interacting with plant AGO1 and that perturbation of their farnesylation was causing changes in AGO1 association with membranes and with membrane-bound polysomes, although the loss of J2 and J3 farnesylation did not impair AGO1 miRNA loading (74). In our case, we qualitatively observed GFP::ALG-1 localization after DNJ-12 depletion and did not detect any gross mislocalization of ALG-1 (Figure S7), suggesting that the mechanism we identified is probably independent of farnesylation and acting upstream in the miRNA pathway before mRNA target binding.

When we performed an in silico analysis to identify any features present in the N-terminal region of ALG-1 using the MobiDB database (75), we noticed that this unique region contains intrinsically disordered residues. A subpart of a similar partially unstructured region to the one predicted in ALG-1 was recently named ‘’the beam’’ in human AGO2, which is predicted to stabilize AGO by interacting with N, L1, L2 and PIWI domains (4). Also, a subpart of *A. thaliana* AGO1 N-terminal domain was defined as the N-terminal coil and is thought to be important for AGO function (76) and interaction with regulatory factors (77). Here, we propose that through its interaction with the specific N-terminal region of ALG-1, DNJ-12 can modulate ALG-1 interaction with the HSP70 system, induce a remodelling of ALG-1 and consequently regulate the loading of ALG-1 by miRNAs (Figure 6). Strikingly, a recent study in *A. thaliana* demonstrates that plant AGO1 contains an unstructured N-terminal region (named NTE for N-terminal extension in this study) important for plant viability as well as miRNA stability and loading. (78). This new observation in plants strengthens our ideas that the N-terminal region of AGO is crucial for miRNA loading, and the mechanism we uncovered is likely widespread across different species.

Our data show that the co-chaperone DNJ-12 can interact with ALG-1 N-terminal region and help connecting AGO to the HSP70 chaperone HSP-1, to load several miRNAs onto the proper AGO in animals. According to the model, it is likely that depletion of HSP70 would result in miRNA defects similar to those seen with DNJ-12 depletion. We attempted to create a worm strain that included an AID-tagged version of HSP-1, along with a construct expressing TIR1 specifically in the seam cells because, as for *dnj-12*, *hsp-1* is essential in animals (79). Although we successfully inserted the AID tag at the N-terminus and internally of the hsp-1 gene, the insertions altered HSP-1 function as we could not isolate viable animals even in the absence of IAA. The fact that we cannot experimentally test the contribution of HSP70 in this process represents a limitation of our study.

The role of the HSP70 (and HSP90) machinery was recently examined in the loading of miRNA duplex onto AGO10 in *Arabidopsis Thaliana* (80). The authors showed that the combination of HSP70 and HSP40 (DnaJ2) increases the loading of miRNA duplexes in AGO10. Interestingly, it was also demonstrated that the HSP70 machinery grants additional loading specificity to AGO10, allowing it to load miR166/165 which are essential for plant development (81). This selection mechanism relies on a central loop formed by the 714–720 residues in AGO10 (80). This loop differs from the one in human AGO2, explaining why this AGO does not display such a strong specificity in duplex loading. When we aligned the sequences of the loops of plant AGO10, AGO2 and ALG-1, we observed that ALG-1 and AGO2 have similar residues, suggesting that the HSP70 machinery, via DNJ-12, is modulating miRNA loading onto ALG-1 through another mechanism (Figure S8A).

Our study uncovered a mechanism in which DNJ-12 modulates ALG-1 loading by miRNAs. We identified a subset of miRNAs that are less loaded onto ALG-1 after DNJ-12 depletion using our small RNA sequencing data. Furthermore, we noticed that when a guide miRNA is downregulated, its complementary star strand is usually also downregulated (Figure 5C), suggesting a model in which ALG-1 fails to load specific miRNA duplexes in the absence of DNJ-12, leading to their degradation. Conversely, in cases where star strands have higher expression after DNJ-12 depletion, we do not detect an increase in the complementary guide strand, suggesting that these star strands might not be ejected appropriately during the miRISC assembly (Figure 5D). Interestingly, some miRNAs were upregulated after tissue-specific depletion of DNJ-12, notably in the seam cells (Figure 4B). Since alae do not form upon DNJ-12 depletion, it is plausible that the seam cells do not terminally differentiate, potentially explaining the change in the miRNA profile. Furthermore, perturbation in AGO homeostasis can cause upregulation of some miRNAs, as seen in the *alg-1* null allele (61). Intriguingly, we do not detect any changes in the level of miR-228, even though miR-228-dependent target repression is altered after DNJ-12 depletion. This discrepancy can be explained by the fact that we monitored the effect of miR-228 repression in specific cells (intestinal cells) and quantified miRNA levels from whole animals. Therefore, the miR-228 level monitored by small RNAs sequencing likely does not reflect its level found only in those specific cells.

From our data, it is interesting to note that DNJ-12 depletion alters only the level of a subset of miRNAs. In constrast, the loss of ALG-1 in animals causes a widespread decrease of miRNAs (61). We believe this difference in the number of downregulated miRNAs can result from several factors. First, this may suggest that only a subset of miRNAs requires the co-chaperone DNJ-12 to be loaded onto ALG-1. It could be possible that structural features of some miRNA duplexes or precursors make them more challenging to be loaded into ALG-1 protein and thus require the co-chaperone/chaperone system to keep this Argonaute in an open conformation for a longer time to permit miRNA loading, as suggested by the Tomari lab (29). It is also possible that due to the extreme stability of some miRISCs (half-lives more than 24h (82)), we are not detecting any effect on some miRNA levels that are loaded at an early stage of the animal development upon DNJ-12 depletion in contrast to alg-1 null animals (As ubiquitous depletion of DNJ-12 leads to sterility, we cannot perform DNJ-12 depletion over generations and thus survey its role in miRNA loading at earlier periods of animal development).

Identifying DNJ-12 as a specific interactor of ALG-1 but not ALG-2 suggests that both proteins, although highly homologous (39), can interact with distinct binding partners through their distinctive regions in the N-terminal domain. Even though ALG-1 and ALG-2 expression patterns mostly overlap spatiotemporally (39, 61) and DNJ-12 is expressed in multiple tissues in *C. elegans* (83, 84), we can envision that the identity of the proteins interacting specifically with those two miRNA-specific Argonautes could explain their partial functional redundancies observed during *C. elegans* development (39, 40, 68, 85). Interestingly, HDJ-2 (*dnj-12* ortholog) interacts with mammalian AGO2, suggesting this chaperone-dependent mechanism might be conserved (21). Furthermore, human AGOs (AGO1, AGO2, AGO3 and AGO4) possess highly divergent N-terminal regions (Figure S8B), indicating that differential interactions may also regulate AGO in humans. Additional studies will be required to assess the conservation of HSP40 interactions with AGOs and their implication on small RNA loading in other species.

## DATA AVAILABILITY

The data underlying this article are available in NCBI’s Gene Expression Omnibus at https://www.ncbi.nlm.nih.gov/geo/ and can be accessed with accession number GSE241034.

## AUTHOR CONTRIBUTIONS

Conceptualization, P-M.F. and M.J.S.; Methodology, P-M.F. and M.J.S.; Investigation, P-M.F., G.J. and I.B.; Writing – Original Draft, P-M.F.; Writing – Review & Editing, P-M.F., G.J. and M.J.S.; Funding Acquisition, M.J.S.; Supervision, M.J.S.

## ACKNOWLEDGEMENTS

We want to thank Dr. Kotaro Nakanishi, Gabrielle St-Onge and the lab members for insightful comments on the manuscript. We also want to thank François Houle, Carl St-Pierre and Laure Villoing for their technical help during the project, and Dr. Weifeng Gu and Vaishnav Vasudevan for their help with small RNA sequencing analysis. We also thank Jordan D. Ward for his technical help with the AID strains. Some strains were provided by the CGC, funded by the NIH Office of Research Infrastructure Programs (P40 OD010440). Pierre-Marc Frédérick received scholarships from Faculté de médecine de l’Université Laval (M.Sc.), Fonds de recherche du Québec-Santé (M.Sc. and Ph.D.) and Natural Sciences and Engineering Research Council (Alexander-Graham-Bell Ph.D.).

## FUNDING

This work was supported by The Canadian Institutes of Health Research (CIHR).

## CONFLICT OF INTEREST

None declared.

## Notes

### Competing Interest Statement

The authors have declared no competing interest.

