## Supplementary Data for "Interaction between a J-domain co-chaperone and a specific Argonaute protein contributes to microRNA function in animals"

### SUPPLEMENTARY FIGURES LEGENDS

#### Figure S1. Homology between the nematode miRNA-specific Argonautes ALG-1 and ALG-2

**(A**) Alignment of the N-terminal regions of ALG-1 (G5EGR6) and ALG-2 (Q86B39). The sequence used to perform the yeast two-hybrid screen is highlighted in yellow. The protein sequences used for alignment are indicated by the Uniprot identifier number.

#### Figure S2. Tissue-specific depletion of DNJ-12 using the AID system

**(A)** Confocal images (merged images of DIC and BFP on a black background) of MJS418 worms grown at 20°C on either EtOH 0.4% or IAA 4mM OP50 plates. Pictures were taken at the late L4 stage. The scale bar represents 20µm. **(B)** Confocal images (merged images of DIC and BFP) of MJS444 and MJS469 worms grown at 20°C on either EtOH 0.4% or IAA 4mM OP50 plates. Pictures were taken at the late L4 stage. The scale bar represents 20µm. **(C)** Western blot of worms boiled in Laemmli buffer (MJS418) after 180 minutes of IAA 4mM treatment and detection of ALG-1, AID::DNJ-12::HA and ACTIN. Worms were grown for 55h at 20°C, then shifted on 4mM IAA small plates for the indicated time. (Cropped version of Figure 3B.) **(D)** Representative images of normal worms (WT) and the vulval bursting phenotype (Burst).

#### Figure S3. Characterization of the effect of IAA treatment, DNJ-12 depletion and DNJ-12 endogenous tagging.

**(A)** (Left) Representative images (GFP) of L4 MJS482 worms (expressing the miR-228 sensor reporter, AID::DNJ-12::HA and the ubiquitously expressed TIR1) grown on either EtOH 0.4% or IAA 4mM plates for their whole post-embryonic development at 20°C. The scale bars represent 20µm. (Right) Representative images (GFP) of L4 MJS488 worms (expressing the miR-228 sensor reporter and the ubiquitously expressed TIR1) grown on either EtOH 0.4% or IAA 4mM plates for their whole post-embryonic development at 20°C. The scale bars represent 20µm. **(B)** (Left) Representative images (GFP) of L4 MJS495 worms (expressing the miR-228 mutated sensor reporter, AID::DNJ-12::HA and the ubiquitously expressed TIR1) grown on either EtOH 0.4% or IAA 4mM plates for their whole post-embryonic development at 20°C. The scale bars represent 20µm.

(Right) Quantification of the GFP signal in arbitrary units of three biological replicates of auxin treatments. In each biological replicate, each dot represents the mean of 4 GFP cells on one single worm. The horizontal bar represents the mean. p-value was calculated using a two-tailed unpaired t-test. **** represents $p \leq 0.0001$. **(C)** Quantification of the bursting phenotype at the adult stage of worms grown on OP50 plate at 20°C or 25°C. **(D)** Quantification of target mRNAs (li*n-41, daf-12, hbl-1 and lin-28*) expression measured by RT-qPCR. RNA was purified from young adult worms grown on EtOH 0.4% or IAA 4mM plates for their whole development at 20°C. The tba-1 mRNA was used as an endogenous control to obtain ΔΔCT values. The error bars indicate the 95% confidence interval. *p*-values were calculated using a two-tailed unpaired t-test. ns represents $p > 0.05$, * represents $p \leq 0.05$ and ** represents $p \leq 0.01$.

**Figure S4. Characterization of the interaction between ALG-1 and DNJ-12**

Pulldown of proteins associated with let-7 in extracts of worms expressing DNJ-12::HA. Proteins associated with let-7 were pulled down using a 2′-*O*-Methyl modified oligonucleotides oligonucleotide fully complementary to let-7. A similar oligonucleotide complementary to the luciferase sequence was used as a control. The pulled-down proteins were analyzed by western blot to detect the presence of ALG-1 and DNJ-12::HA. 50ug and 2mg of total lysate were used for input and pulldown, respectively.

**Figure S5. Additional characterization of total downregulated miRNAs after DNJ-12 tissue-specific depletions**

**(A)** Principal component analysis plot showing the first two principal components of reduced RNA sequencing data (RPM + 1). The dots with the same color represent biological replicates for the given conditions (tissue of depletion and EtOH or IAA treatment) in the legend. **(B)** miRNA sequencing data (total small RNAs) plotted according to IAA-mediated DNJ-12 depletion in the seam cells, the pharynx or ubiquitously. The raw reads of three biological replicates were individually normalized on the total gene expression reads (geex) minus the structural reads (struct) and converted to RPM. The Y axis represents the average $\log_2$ RPM of IAA-treated worms and the X axis represents the average $\log_2$ RPM of EtOH-treated worms. Each dot represents one miRNA. The downregulated miRNAs and their $\Delta\log_2$(IAA RPM – EtOH RPM) values are listed on the right.

**Figure S6. Additional characterization of IPed downregulated miRNAs after DNJ-12 tissue-specific depletion**

**(A**) Relative quantification of other small RNA types. We normalized the levels of 21U RNAs (PRG-1) and 26G RNAs (ALG-3/ALG-4) on the total gene expression reads (geex) minus the structural reads (struct). Relative values of every replicate were plotted. The error bars represent standard deviation.

**(B)** miRNA sequencing data (IPed small RNAs) plotted according to IAA-mediated DNJ-12 depletion ubiquitously. The raw reads of three biological replicates were individually normalized on the level of ALG-1 protein immunoprecipitated. The Y axis represents the average $\log_2$ values of IAA-treated worms and the X axis represents the average $\log_2$ values of EtOH-treated worms. Each dot represents one miRNA. The downregulated miRNAs and their $\Delta\log_2$(IAA – EtOH) values are listed on the right.

**Figure S7. DNJ-12 depletion in the pharynx does not lead to mislocalization of ALG-1**

Representative confocal images of ALG-1 localization (BFP and GFP merge) of young adults MJS487 worms (expressing GFP::AGL-1, AID::DNJ-12::HA and seam cell TIR1) grown on either EtOH 0.4% or IAA 4mM plates for their whole post-embryonic development at 20°C. The scale bars represent 10µm.

**Figure S8. Homology between central loop of plant AGO10, worm ALG-1 and human AGO2, and N-terminal of human AGO1-4**

**(A)** Alignment of the central loop (714-720) of AGO10 (*A. thaliana*), ALG-1 (*C. elegans*) and AGO2 (*H. sapiens*).

**(B)** Alignment of the N-terminal region of human AGO1 (Q9UL18), AGO2 (Q9UKV8), AGO3 (Q9H9G7) and AGO4 (Q9HCK5). The protein sequences used for alignment are indicated by the Uniprot identifier number.

**SUPPLEMENTARY TABLES (EXCEL FILE)**

**Table S1:** List of nematode strains used in this study

**Table S2:** List of oligos used in this study

**Table S3**: Western blot quantifications

**Table S4:** miR-228 reporters quantification

**Table S5:** Box B / ΔN GFP reporter quantification

**Table S6:** Small RNA sequencing data (Seam cell promoter total)

**Table S7:** Small RNA sequencing data (Pharynx promoter total)

**Table S8:** Small RNA sequencing data (Ubiquitous promoter total)

**Table S9:** Small RNA sequencing data (input downregulated and upregulated microRNA)

**Table S10:** Small RNA sequencing data (Ubiquitous promoter IP)

**Table S11:** Small RNA sequencing data (IP downregulated and upregulated microRNA)

**Table S12:** Small RNA sequencing data (ubiquitous promoter 21U and 26G RNAs)

N-terminal

```
ALG-1-G5EGR6      MSGGPQYLPGVMNSTIQQQPQSATSSFLPSGPISSTSTSSQVVPTSGATQQPPFPSAQAA  60
ALG-2-Q86B39      ------------------------------------------------------------  0


ALG-1-G5EGR6      ASTALQNDLEEIFNSPPTQPQTFSDVPQRQAGSLAPGVPIGNTSVS-IGEPANTLGGGLP  119
ALG-2-Q86B39      ----------------------------------MPGDSLTSSSFMPDGGAETSSSSQLG  26
                                                   **  : .:*.   *   .: .. *

ALG-1-G5EGR6      GGAPGQLPGGNQSGIQFQCPRRPNHGVEGRSILLRANHFAVRIPGGTIQHYQVDVTPDKC  179
ALG-2-Q86B39      GSAHGAIGTKPDAGVQFQCPVRPNHGVEGRSILLRANHFAVRIPGGSVQHYQIDVFPDKC  86
                  *.* * :    ::*:***** *************************::****:** ****
```

Frédérick et al., Figure S1

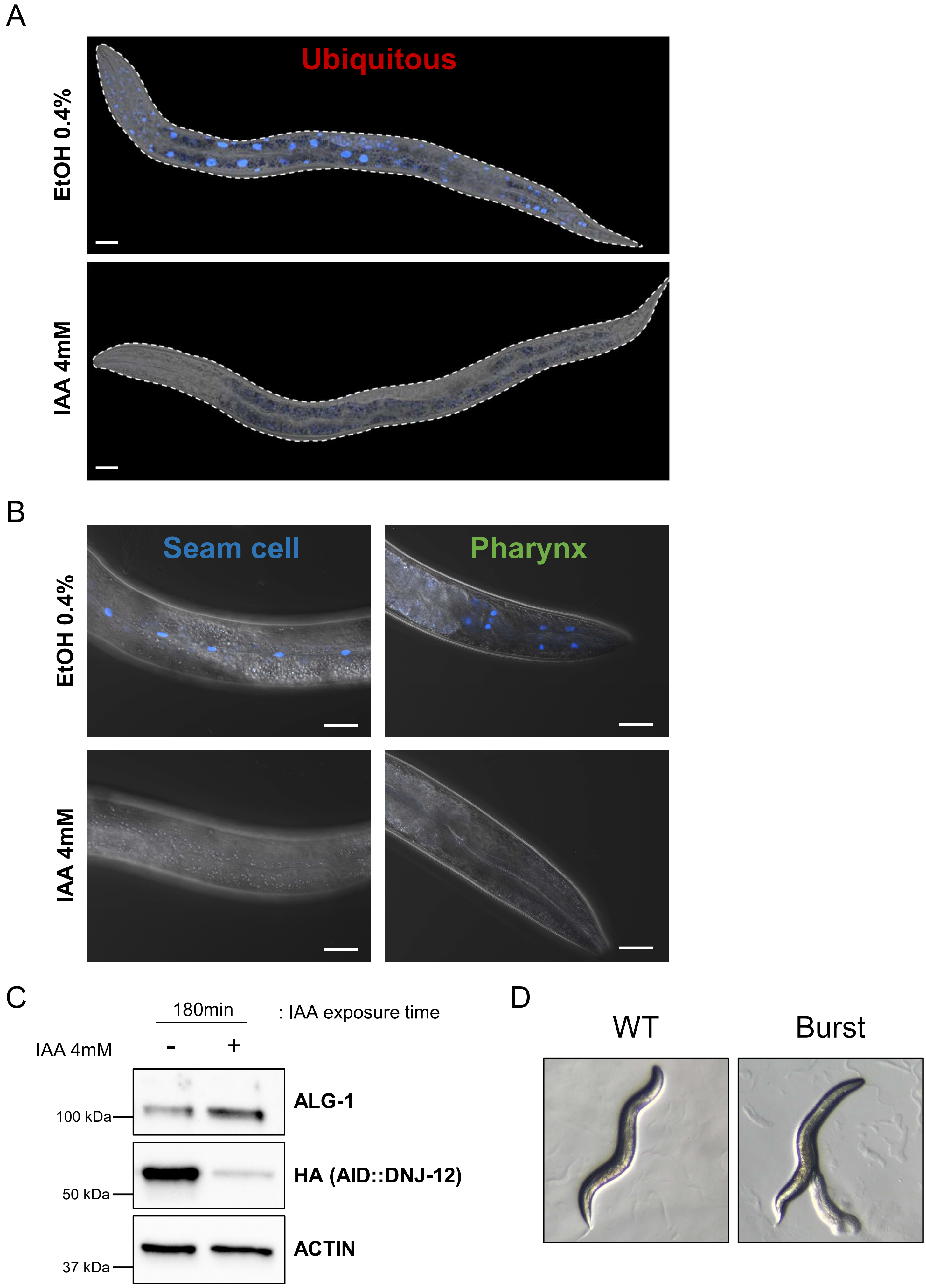


Frédérick et al., Figure S2

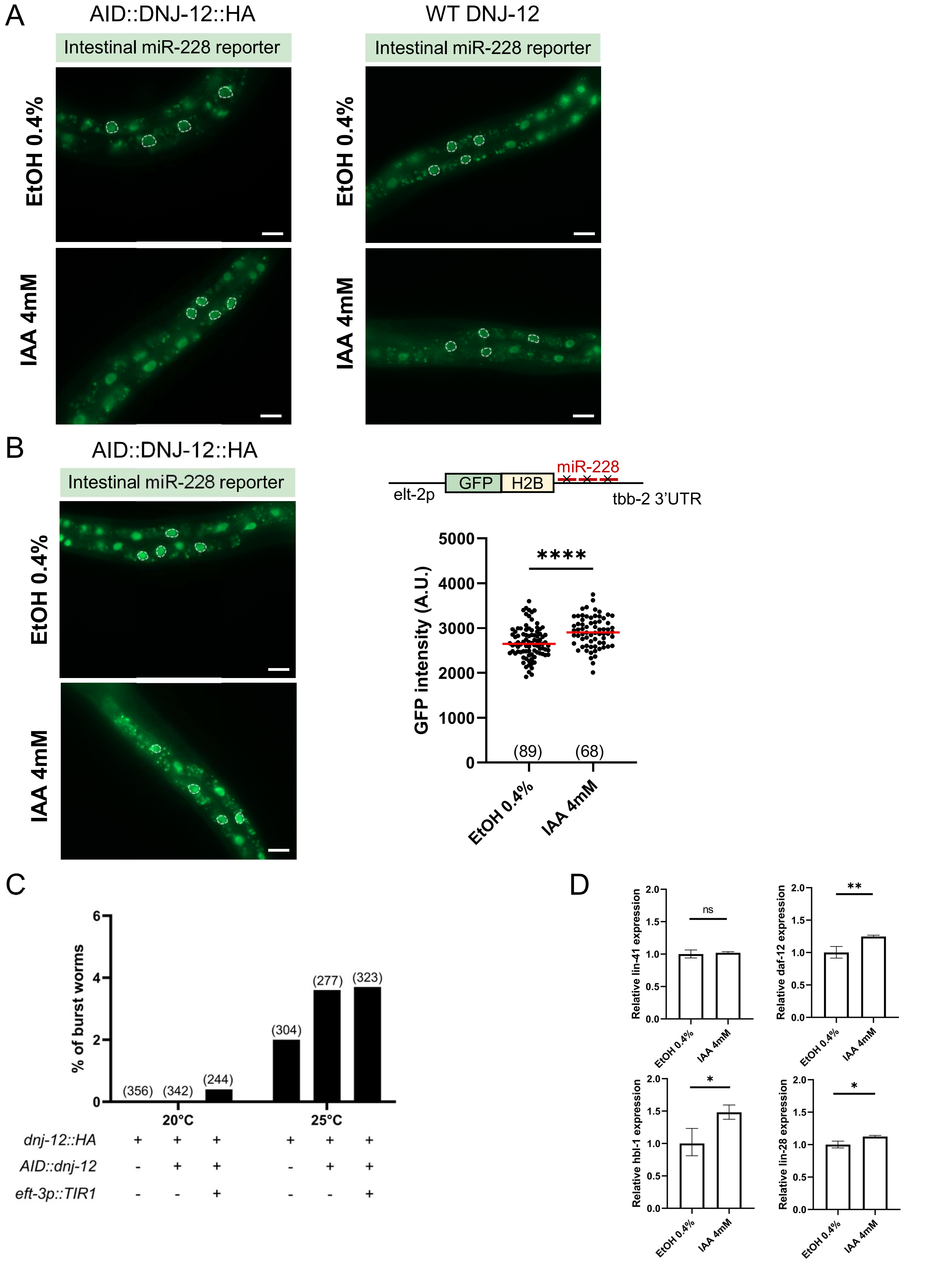


Frédérick et al., Figure S3

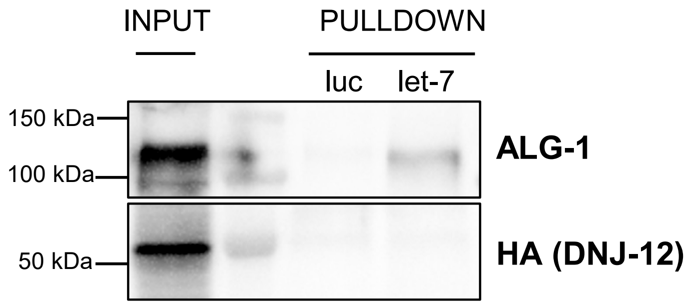


Frédérick et al., Figure S4

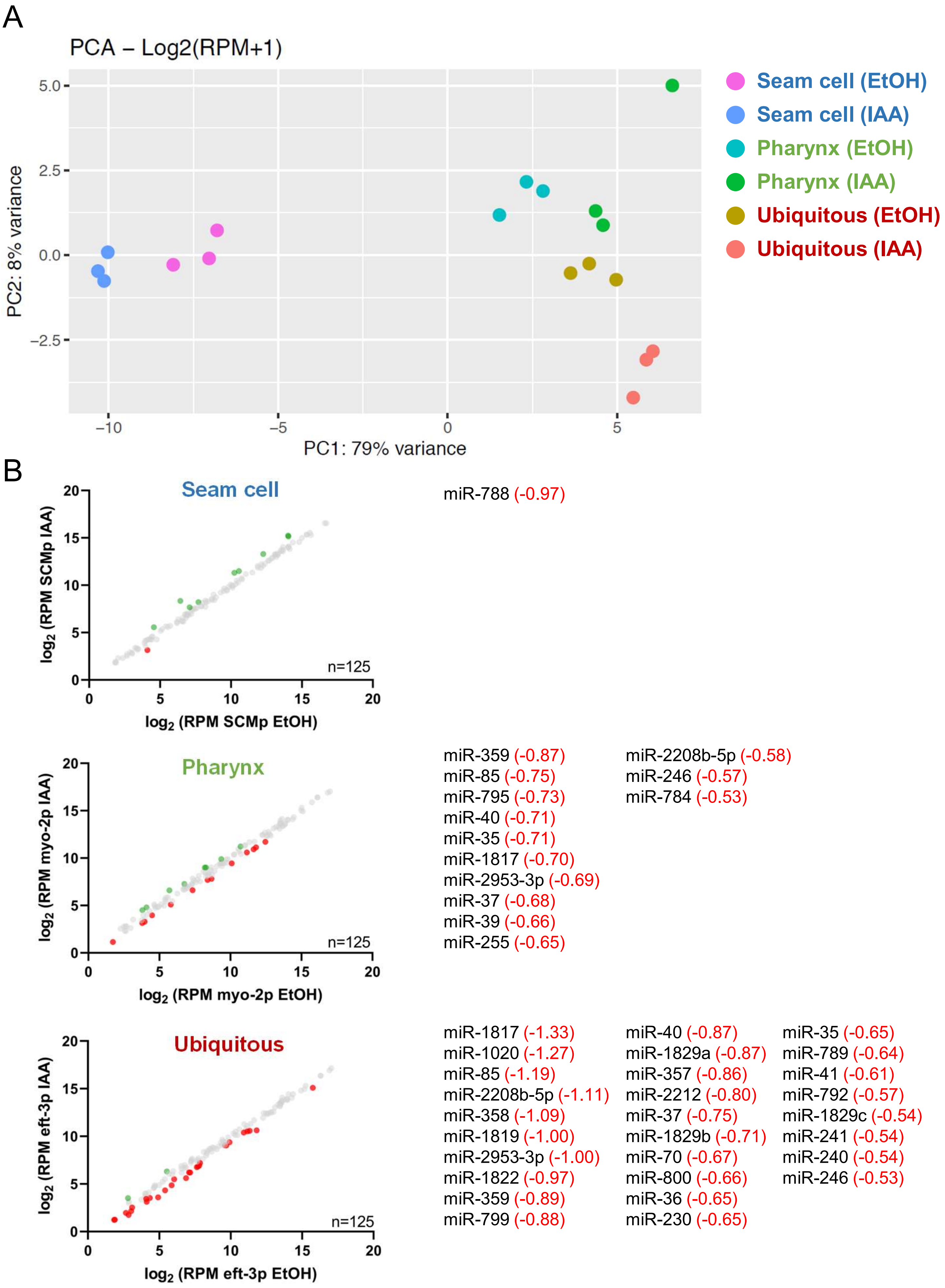


Frédérick et al., Figure S5

A

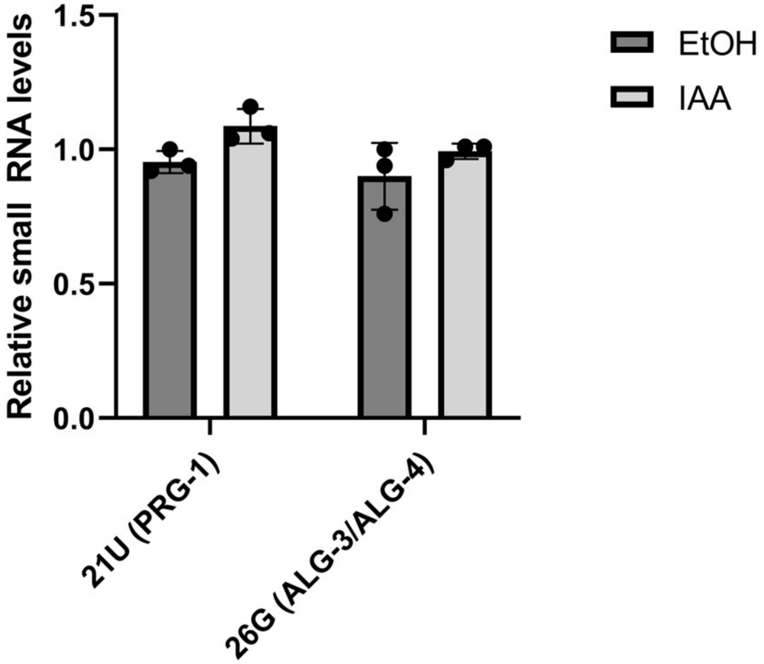


B

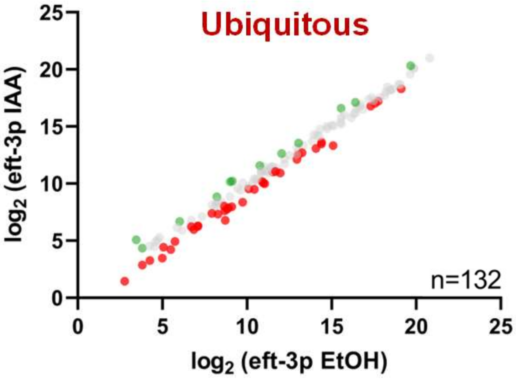


| | | | |
|---|---|---|---|
| miR-1817 (-1.93) | miR-786 (-1.04) | miR-800 (-0.85) | miR-227 (-0.57) |
| miR-85 (-1.76) | miR-1020 (-1.03) | miR-241 (-0.84) | miR-39 (-0.56) |
| miR-2217 (-1.52) | miR-246 (-1.00) | miR-70 (-0.81) | let-7 (-0.54) |
| miR-1829a (-1.39) | miR-41 (-0.99) | miR-2208b-5p (-0.81) | miR-38 (-0.54) |
| miR-2209a (-1.33) | miR-1819 (-0.95) | miR-2212 (-0.79) | miR-65 (-0.53) |
| miR-1829b (-1.27) | miR-1832 (-0.95) | miR-37 (-0.79) | miR-252 (-0.53) |
| miR-799 (-1.14) | miR-359 (-0.94) | miR-357 (-0.73) | miR-42 (-0.52) |
| miR-1822 (-1.09) | miR-35 (-0.93) | miR-2208b-3p (-0.63) | miR-237 (-0.52) |
| miR-40 (-1.06) | miR-36 (-0.92) | miR-795 (-0.63) | miR-784 (-0.50) |
| miR-358 (-1.05) | miR-2953-3p (-0.89) | miR-1829c (-0.61) | |

Frédérick et al., Figure S6

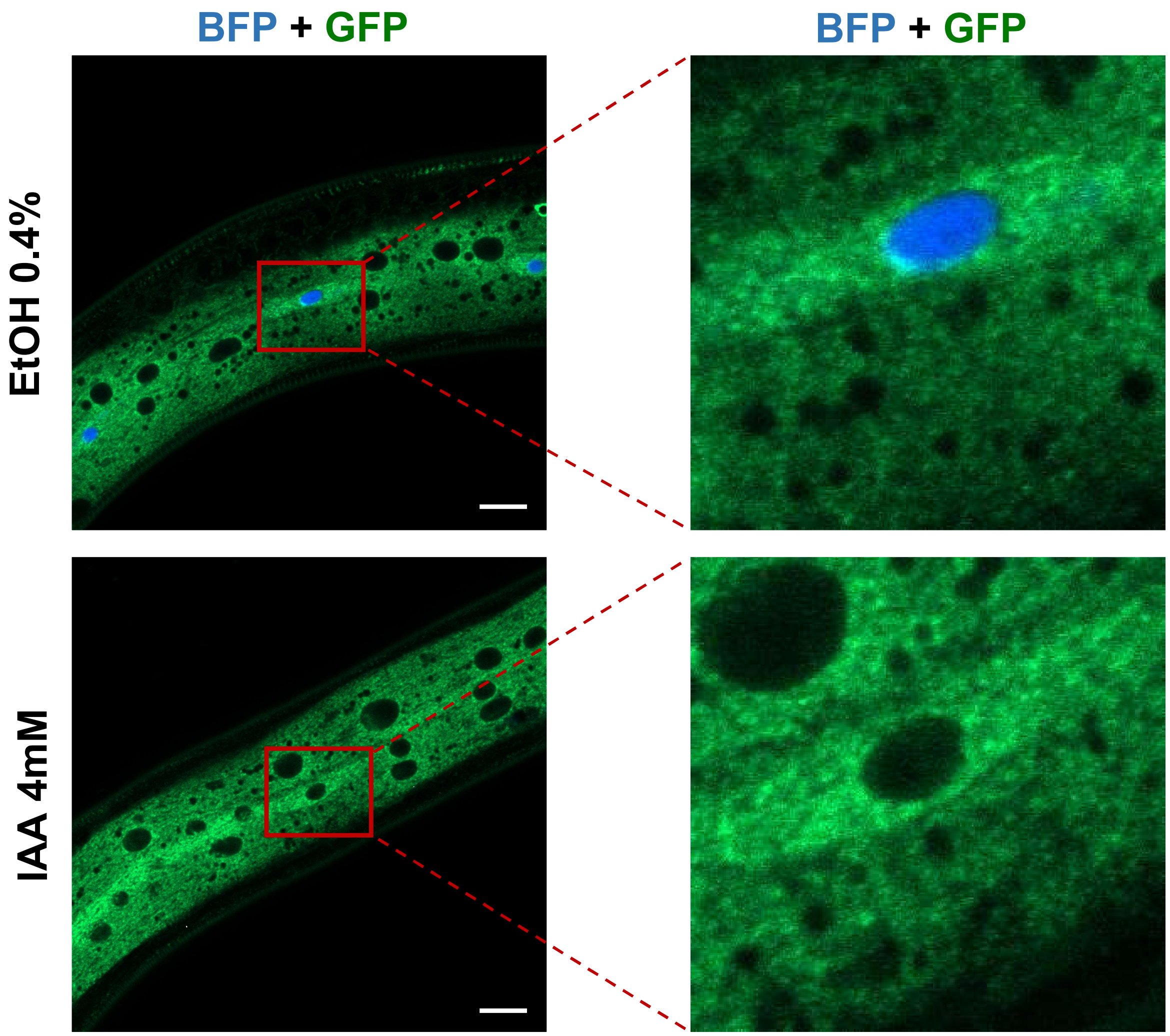


Frédérick et al., Figure S7

A

```
                    714-720
AGO10      DVTHPENGEESSPSIAAVVAS
ALG-1      DITHPPAGDSRKPSIAAVVGS
AGO2       DVTHPPAGDGKKPSIAAVVGS
           *:***  *:  .*******.*
```

B

N-terminal

```
AGO1-Q9UL18      MEAGPSGAAAGAYLP---PLQQVFQAPRRPGIGTVGKPIKLLANYFEVDIPKIDVYHYEV   57
AGO2-Q9UKV8      -MYSGAGPALAPPAPPPPIQGYAFKPPPRPDFGTSGRTIKLQANFFEMDIPKIDIYHYEL   59
AGO3-Q9H9G7      MEIGSAGPAG---------AQPLLMVPRRPGYGTMGKPIKLLANCFQVEIPKIDVYLYEV   51
AGO4-Q9HCK5      MEALGP--------G---PPASLFQPPRRPGLGTVGKPIRLLANHFQVQIPKIDVYHYDV   49
                                        :  * **. ** *: *:* ** *:::*****:* *::

AGO1-Q9UL18      DIKPDKCPRRVNREVVEYMVQHFKPQIFGDRKPVYDGKKNIYTVTALPIGNERVDFEVTI   117
AGO2-Q9UKV8      DIKPEKCPRRVNREIVEHMVQHFKTQIFGDRKPVFDGRKNLYTAMPLPIGRDKVELEVTL   119
AGO3-Q9H9G7      DIKPDKCPRRVNREVVDSMVQHFKVTIFGDRRPVYDGKRSLYTANPLPVATTGVDLDVTL   111
AGO4-Q9HCK5      DIKPEKRPRRVNREVVDTMVRHFKMQIFGDRQPGYDGKRNMYTAHPLPIGRDRVDMEVTL   109
                 ****:* *******:*: **:***  *****:* :**::.:**.  **:.   *:::**:

AGO1-Q9UL18      PGEG-KDRIFKVSIKWLAIVSWRMLHEALVSGQIPVPLE--------SVQALDVAMRHLA   168
AGO2-Q9UKV8      PGEG-KDRIFKVSIKWVSCVSLQALHDALSGRLPSVPFE--------TIQALDVVMRHLP   170
AGO3-Q9H9G7      PGEGGKDRPFKVSIKFVSRVSWHLLHEVLTGRTLPEPLELDKPISTNPVHAVDVVLRHLP   171
AGO4-Q9HCK5      PGEG-KDQTFKVSVQWVSVVSLQLLLEALAGHLNEVP--------DDSVQALDVITRHLP   160
                 **** **: ****::::: ** : * :.* .     *           ::*:**  ***
```

Frédérick et al., Figure S8
